# The effect of urbanisation and local environmental heterogeneity on phenotypic variability of a tropical treefrog

**DOI:** 10.1101/2024.11.28.625871

**Authors:** Marcos R. Severgnini, Diogo B. Provete

## Abstract

1. Urbanisation reduces species richness and change community composition. However, little is known on how the phenotype of organisms with low dispersal ability respond to environmental changes associated with urbanisation in fast urbanizing centres, such as those in the Global South.

2. Here, we tested how urbanisation rate, local environmental heterogeneity, land surface temperature, and spatial gradients affect phenotypic traits associated with dispersal, resource acquisition, and performance, namely: body size, head shape, and leg length of the Dwarf Treefrog (*Dendropsophus nanus*) using a space-for-time substitution approach.

3. We took linear measurements from 768 individuals in 21 ponds along an urban gradient in central Brazil. We also measured local environmental variables and summarized them using Hill-Smith Principal Component Analysis. The spatial arrangement of ponds at multiple scales was described using Moran Eigenvector Maps. Those variables were then entered into a Structural Equation Model to test their direct and indirect effects on the mean and coefficient of variation (CV) of phenotypic traits. Additionally, we calculated the Scaled Mass Index as a proxy for fitness and estimated the adaptive landscape for body size, size-free leg length, and head shape. We also tested for spatial autocorrelation in traits.

4. Body size decreased from the periphery to the urban centre, whereas CV of body size and head shape had the opposite pattern. Body size increased, whereas CV of body size and head shape decreased in man-made ponds. The CV of leg length decreased with increasing land surface temperature. The remaining traits were not affected by any predictor variable. None of the traits were spatially autocorrelated. Both body size and head shape were under weak directional selection, but in opposite directions.

5. Our results suggest that the lack of a clear spatial variation in phenotypic traits can be due to a weak selection, due to a recent, although intense, urbanisation process. In conclusion, eco-evolutionary dynamics in tropical cities seem to have a different pace compared to temperate ones. Our results can contribute to building urban ecological theory that explicitly includes city age, their development, growth rate, and history.

## 1. INTRODUCTION

Novel ecosystems, such as cities can be challenging to resident species because their altered environment to benefit humans (e.g., artificial light at night, anthropogenic noise, chemical stressors) might not be suitable for all species. Urbanisation has negatively been impacting biodiversity and altering the pace of natural selection around the globe (Alberti, 2015; Hendry et al., 2008). Built environments promote habitat fragmentation and isolation, reduce gene flow, and create small populations more prone to genetic drift (Miles et al., 2019; Rivkin et al., 2019). Species may respond to these novel ecological conditions by changing either their morphology, physiology, and behaviour, or by exaptation (Winchell et al., 2023). For example, the urban heat island should impose strong metabolic costs to organisms, leading to a decrease in body size (Merckx et al. 2018). Indeed, some studies (e.g., Jennette et al., 2019) have found smaller body sized in urban populations of frogs when controlled for age. However, larger body sizes are usually associated with increased fitness (Smith & Belk, 2018). Consequently, maladapted phenotypes may be fixed in urban populations, decreasing survival and reproduction of individuals (Urban, 2011; Brady et al., 2019a).

Recent studies found that urbanisation changes morphological traits of birds, reptiles, and mammals (Alberti et al., 2017; Fugère & Hendry, 2018). However, organisms with high dispersal ability (e.g., birds) can still maintain moderate gene flow (Miles et al., 2019) and are more likely to adapt to urban environments (Schmidt et al., 2020). Conversely, the effects of urbanisation on organisms with low dispersal ability, like frogs, are expected to be more pronounced (Khimoun et al., 2020), because dispersal limitation reduces gene flow and population size. Dispersal can also mediate how individuals assess habitats that maximize their performance (Edelaar et al., 2008), resulting in higher phenotypic variation in more heterogeneous habitats (Thompson et al. 2022). Yet, little is known about how phenotype-environment relationships respond to urbanisation, especially in the Global South (Severgnini et al. *in press*), which has been experiencing a fast and recent urbanisation process (Myers, 2021; Shackleton et al., 2021). Frogs are ideal organisms to investigate this question (Hamer & McDonnell, 2008), since they have low dispersal ability and plastic phenotypes modulated by the environment (Levis et al., 2018).

Although several studies have evaluated how phenotypic traits are affected by urban ecosystems (Callaghan et al., 2021; Jennette et al., 2019; Komine et al., 2022), most of them focused on trait mean and did not incorporate phenotypic variance (but see Thompson et al., 2022). While the phenotypic mean can reveal if a trait has already been or will be shaped by natural selection, phenotypic variation can help identify if some trait values in a population have a better fit to the environment (Edelaar et al. 2008; Sanderson et al. 2023), and why they evolve more rapidly (Des Roches et al., 2018). Intraspecific variability modulates several ecological and evolutionary processes (Moran et al., 2016 Sanderson et al. 2023). It can be caused by different processes, such as genetic drift, mutation, phenotypic plasticity, or relaxed natural selection (Sanderson et al. 2023). Phenotypic variance, especially intraspecific variability, can provide a different perspective for urban evolutionary studies. For example, phenotypic variance of two bird species increased, but had lower means, in urban sites (Thompson et al., 2022) compared to their forest counterparts, likely due to disruptions in developmental pathways. However, dispersal limited groups, whose distribution is constrained to the surroundings of water bodies, might experience a decrease in phenotypic variation due to lower population sizes as a result of habitat connectivity loss (Thompson et al., 2022). In fact, a previous study across two urbanised landscapes found gene flow happened up to 6 km in a frog (Homola et al. 2019).

Previous studies have found a strong effect of urban ecosystems on the tempo and mode of phenotypic evolution (Diamond & Martin, 2021). Urban populations usually experience relaxed selection, but divergence between urban and non-urban populations has often led to nonparallel evolution (Rivkin et al., 2019; Thompson et al., 2022). However, few studies have evaluated how selection varies along urbanisation gradients, and none of them focused on semi-aquatic organisms, such as frogs. Thus, an important, but open question (see Thompson et al., 2022) is how the selection on different types of traits differ in strength and mode within cities.

Most urban evolutionary studies have been conducted in the Northern Hemisphere (reviewed in Szulkin et al., 2020). However, urban ecosystems around the globe are not equal. Instead, cities have distinct modes of planning, occupation histories, ages, growth rates, and socio-economic aspects (Myers, 2021; Shackleton et al., 2021). Phenotypic variation can respond in different ways to urban ecosystems, since cities may impose similar, but not equal challenges to species (Thompson et al., 2022; delBarco-Trillo & Putman, 2023). There is also a lack of knowledge on how multiple phenotypic dimensions, such as those associated with dispersal ability, resource acquisition, and performance (McPeek, 2017) change along urban gradients (Thompson et al., 2022). Furthermore, tropical regions have distinct environmental characteristics, such as higher temperature and humidity, as well as higher diversification rates and species richness (Brown, 2014). As a consequence, their regional species pool can respond to urbanisation in contrasting ways, as compared to temperate ones (Shackleton et al., 2021). Although recent, urbanisation in the tropics have been intense (Marcacci et al., 2021), with specially accelerated rates in Latin America and Africa (Myers, 2021). Therefore, understanding how species adapt and thrive in urban environments in the Global South is essential to mitigate negative effects of human impacts (Li et al., 2022).

Here, we tested how urbanisation rate, local environmental heterogeneity, and spatial gradients affect phenotypic traits associated with dispersal, resource acquisition, and jumping performance in the Dwarf Treefrog (*Dendropsophus nanus*) using a space-for-time substitution approach (Wogan & Wang, 2018). This approach assumes that peri-urban environments have changed more rapidly than those occupied by core urban areas. As the urbanisation process intensifies, it creates spatially-structured environments that changed at different times, enabling us to sample populations along space and make inferences about evolution. In this context, we assume frogs living in ponds with high urbanisation rates have been under a strong selective pressure for longer than those in areas that have been urbanized recently. We expect that: (i) both the mean and coefficient of variation (CV) of body size, leg length, and head shape decrease with an increasing urbanisation and increase with local heterogeneity; (ii) mean and CV of body size, leg length, and head shape decrease with land surface temperature; (iii) the adaptive landscape have different peaks for urban and rural populations, with the core urban population having lower fitness than those at less urbanized sites (Figure 1). This is the first study focusing on how both the mean and variance of phenotypes (see Severgnini et al. *in press*) change along urban gradients, while also incorporating spatial structure and indirect relationships between local and landscape variables in a young, tropical city of the Global South.

**Figure 1.**
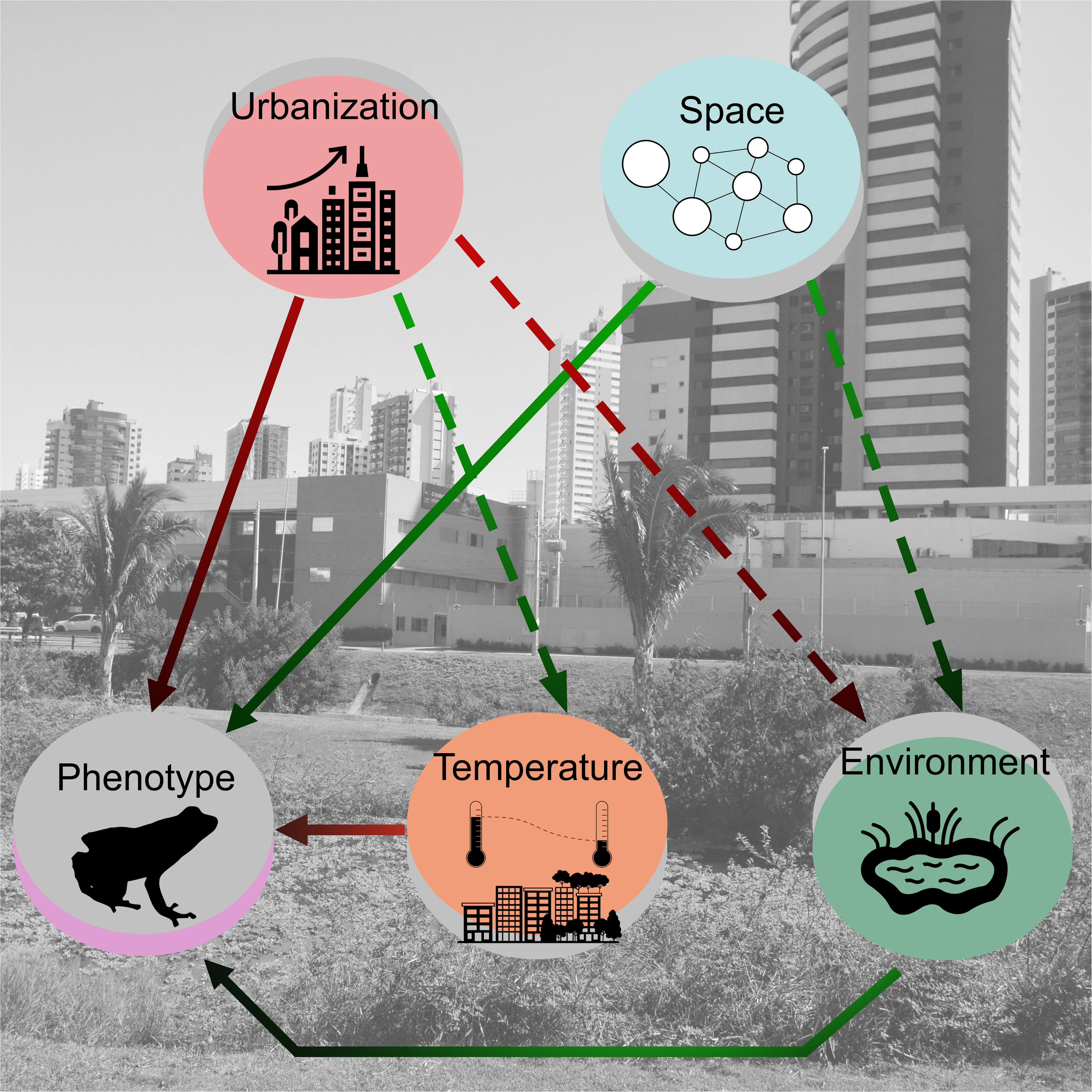
Hypothesis for the study. Green arrows represent positive and red arrows represent negative effects. Dashed arrows represent indirect and continuous arrows represent direct effects. Urbanisation was measured as rate of change from 1985 to 2021. Space represents the spatial arrangement of ponds quantified by Moran’s Eigenvector Maps. The environment was represented by the first axis of PCA summarizing local environmental variables. Temperature is land surface temperature/ urban heat island. Phenotype represents mean or coefficient of variation of phenotypic traits. Frog silhouette is CC-BY from PhyloPic. Environment, urbanization, and temperature silhouettes are CC-BY 3.0 from Noun Project (ponds created by Ircham; urban heat island created by Softscape, and urbanization rate created by WiStudio).

## 2. METHODS

### 2.1 Study site and sampling design

We conducted fieldwork in Campo Grande, Mato Grosso do Sul, central Brazil (Projection: UTM–21S; Coordinates: 751114, 7731659; DATUM=SIRGAS/2000; 529 m a.s.l.; Figure 2). Campo Grande has been founded 127 years ago (Arruda 2006; PLANURB 2019) and has about 897,938 citizens (IBGE, 2023). Although recent, urbanisation has been fast, with the city gaining ∼ 13,000 ha of urban infrastructure from 1985 to 2021 (MRS, pers. obs. based on data from MapBiomas, 2023). The urban area has about 35,903 ha with most human population living in the urban centre (PLANURB, 2019). The climate is Equatorial Savanna or Köppen’s Aw, with dry winters and rainy summers (Kottek et al., 2006). Vegetation is composed mainly by *Cerradão*, *Cerrado*, *Campo Sujo*, and *Vereda* vegetation types (Ribeiro & Walter, 2008).

**Figure 2.**
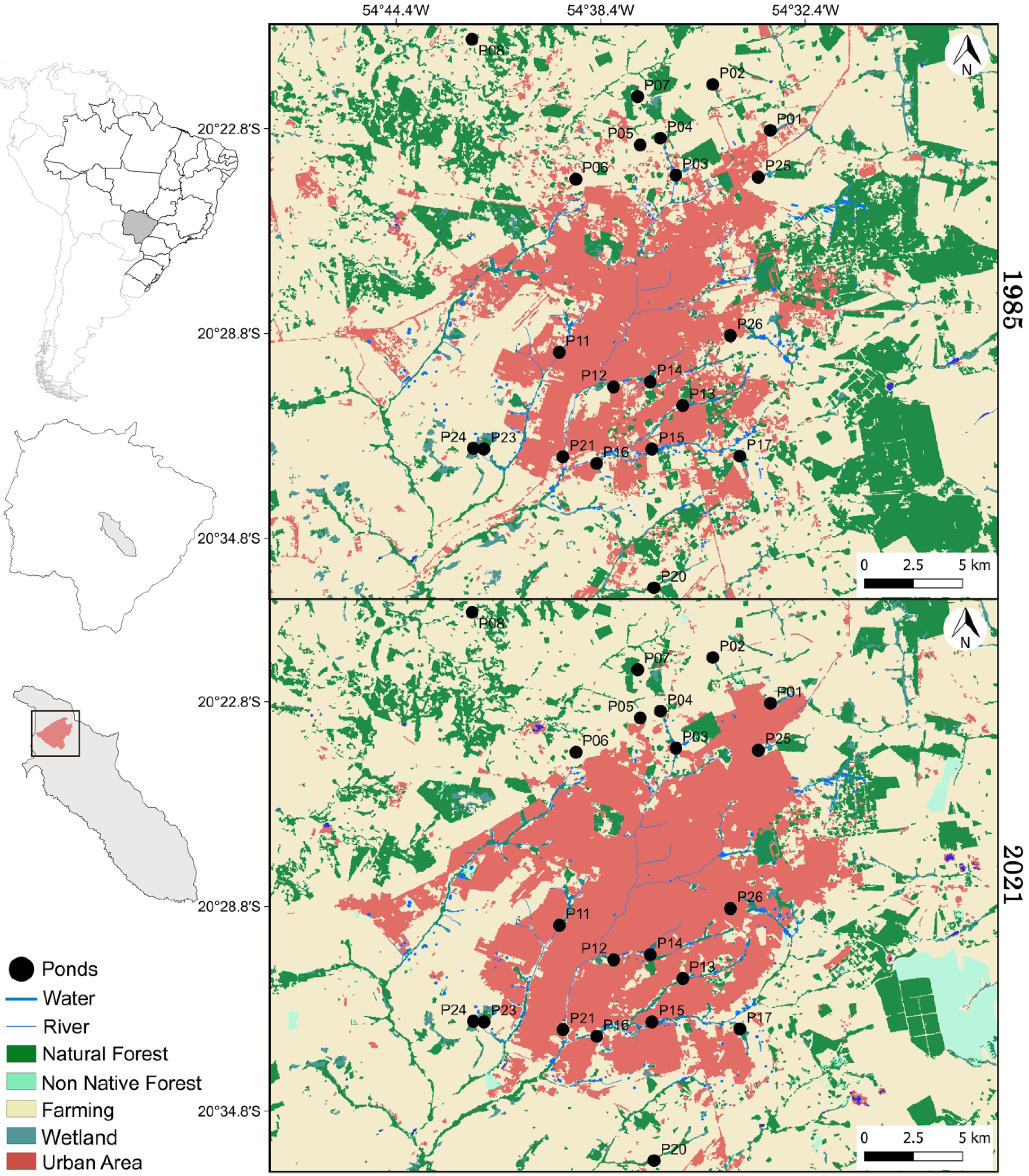
Map of land use and land cover change (1985–2021) showing ponds sampled (black spots) along the rural–urban gradient in Campo Grande, Mato Grosso do Sul, Brazil. Map features extracted from Instituto Brasileiro de Geografia e Estatística (IBGE) and MapBiomas data base 2023; and prepared on QGIS v. 3.22.1.

We sampled adults of the hylid frog *Dendropsophus nanus* in 21 ponds (Figure 2; Figure S1) along an urban gradient through surveys at breeding sites (Scott Jr. et al., 1994) and visual encounter surveys (Crump & Scott Jr., 1994). Field work was conducted between 17:30 and midnight from November 2021 to April 2022 and from November 2022 to January 2023 visiting two ponds per day. We did not visit all ponds in all months. Instead, each pond was visited until we obtained 30 individuals to guarantee equal sampling effort.

### 2.2 Phenotypic traits

We took the following linear measurements in the field using hand digital calliper (MTX 150 mm) to the nearest 0.01 mm: body size (Snout-Vent Length – SVL), head width, head length, and leg length (foot, thigh, and tibiofibula) following Watters et al. (2016). Relative leg length is related to vulnerability to predation (Citadini et al., 2018; Emerson, 1985a), jumping performance, and dispersal ability (Phillips et al., 2006), while head length and width is related to size and variety of feeding resources consumed (Emerson, 1985b; Parmelee, 1999). Body size is related to thermoregulation, dispersal, and desiccation resistance (Wells, 2007). After measurements, every frog was tagged with visible implant elastomer (VIE) and released back.

Prior to analysis, we transformed head width, head length, and leg length to remove the effect of body size by applying a linear regression on each trait as a function of body size and then retained the residuals. To obtain head shape, we took the log-shape ratio (Mosimann, 1970) by computing the geometric mean of head length and head width to obtain their sizes. Then, we divided each value by their geometric means and log-transformed the result (Claude, 2013). Finally, we performed a Principal Component Analysis and took the first axis to represent head shape. PC1 retained 92.5% of the variance and was negatively correlated with head length (-0.74) and width (-0.67). Negative scores along PC1 represent individuals with wider and longer heads, while positive ones represent those with shorter and narrower heads. Analysis was performed in R v. 4.2.3 (R Core Team, 2023).

### 2.3 Fitness proxy

We weighted all individuals using a Pesola (Lightline 10 g #10010) to calculate the Scaled Mass Index (SMI), following Peig & Green (2009). This index relates body mass and size using a Major Axis Regression to obtain a measure of body condition, which was used here as a fitness proxy. Analysis was conducted in lmodel2 R package (Legendre, 2018). The SMI assumes a non-linear growth relationship that removes the effect of body size. This index has been used to access body condition of amphibians, especially in urbanized environments (e.g., Franco-Belussi et al., 2024), since it provides information about animal health status and is positively related with lipid and protein content (MacCracken & Stebbings, 2012). Lipid content in turn is correlated with survival in amphibians (Scott et al. 2007), while body condition also seems directly correlated with survival in a wild toad population (Reading 2007). Field measurements of fitness components are challenging for many taxa, including frogs. We acknowledge that using SMI as a fitness proxy can be problematic (Wilder et al., 2016). However, evidence exists linking lipidic storage to energetic reserves and ultimately to physiological status in frogs. In this context, SMI can be considered a performance index (but see Franklin & Morrissey, 2017), since animals with good body condition that have energetic reserves provided by a past successful feeding are more likely to survive (Jakob et al., 1996; Wells, 2007).

### 2.4 Urbanisation and local environmental heterogeneity

We measured urbanisation as the percentage of building area and roads (Szulkin, Garroway, et al., 2020) in a 500-m buffer around each pond (Figure S2). We also extracted this same variable from raster layers from 1985 to 2021 provided by MapBiomas (2023). To obtain urbanisation rate, we subtracted the current (2021) urban area from that of 1985 and divide it by 36 (number of years in the period). We repeated all analyses using a 1-km buffer and results did not differ. Spatial data handling was conducted in the R packages terra (Hijmans, 2023) and landscapemetrics (Hesselbarth et al., 2019).

To quantify local environmental heterogeneity, we measured the following variables at pond level: pond area (m^2^; using Google Earth®), pond depth (mean of five points per pond; in m), pond hydroperiod (permanent or temporary – binary variable), presence or absence of cattle (binary variable), aquatic predators (vertebrate or both invertebrate and vertebrate – categorical variable), margin profile (excavated, flat or sloping – categorical variable), water temperature (using a thermometer submerged at 1 m for 1 min; in °C), and percentage of floating vegetation (visual estimation %). Ponds with excavated margins were artificial, man-made ponds. Likewise, ponds with vertebrate predators and cattle were more associated with rural environments (Figure S1). Pond margin type limits the amount and kinds of perches available for frogs, influencing their use of the environment (Vasconcelos et al. 2009), likely constraining the range of phenotypic traits associated with each margin type. All variables were later summarized using a Hill-Smith Principal Component Analysis (Hill & Smith, 1976), which allows combining discrete and quantitative variables. The PC1 retained ∼ 63% of the variance in the data. The variables that most contributed to PC1 (correlation > 0.6) were: excavated margin (positive), temporary hydroperiod (negative), invertebrate predators (negative), and cattle presence (negative; Table S1, Figure S3 and S4). Analysis was conducted in the ade4 R package (Dray & Dufour, 2007).

Lastly, we calculated land surface temperature (LST) and urban heat island (UHI; Figure S5). We used the following satellite images: LANDSAT 8 thermal band 10 for 2022 (07 November 2022) and LANDSAT 5 thermal band 6 for 1985 (17 November 1985). All images were obtained from Earth Explorer (United States Geological Survey – USGS: https://earthexplorer.usgs.gov/). The pixels of these images contain solar light variables, such as reflectance and radiance. To extract land surface temperature, we used the equation proposed by Coelho & Correa (2013), which converts digital numbers to solar radiance, radiance to brightness temperature, and then to temperature in Kelvin or Celsius. Then, we used land surface temperature to calculate urban heat island (UHI) (see Ahmed et al., 2013; Faisal et al., 2021) for the whole urban area, as follows:

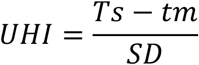

where, *Ts* is land surface temperature; *tm* is the mean of land surface temperature, and SD is its standard deviation. Finally, we built an urban heat island map and calculated maximum, minimum, variance, and mean temperature in 500-m buffers around each pond. Mean land surface temperature was later used as predictor variable in the path analysis. Data handling and extraction were done in QGIS v. 3.22.1 (QGIS.org, 2020).

### 2.5 Building spatial predictors and spatial autocorrelation

To describe the spatial arrangement of ponds at multiple scales, we used Moran’s Eigenvector Maps (MEMs). Firstly, we used the geographical coordinates of ponds to build a spatial neighbourhood network. We tested different types of networks that represent different hypotheses of connections between ponds (i.e., Delaunay triangulation, Gabriel graphs, Minimum spanning trees, and a custom-built connectivity network). Then, we built a spatial weighting matrix in which weighting was a linear function of the inverse of distance. Afterwards, we generated a set of MEMs for each spatial neighbourhood network. Finally, we performed a selection of spatial matrices in the adespatial R package (Dray et al., 2023). We retained the best subset of MEMs that represents the autocorrelation of body size, built from the best spatial neighbourhood, which was selected based on the Akaike’s Information Criterion corrected for small sample sizes (AICc). Then, we computed the corresponding Moran’s *I* statistic of each MEM and calculated its significance using a Monte Carlo permutation test to retain only eigenvectors with positive and significant autocorrelation.

The best spatial model was the custom neighbour network with six positive and significant MEMs (Figure S6). MEM3 had the lowest AICc and was used to represent pond spatial arrangement in the path analysis. Positive scores along MEM3 were arranged in a Northeast-Southwest direction and comprised most of the urbanized ponds (Figure S2 and S6). Finally, to test for spatial autocorrelation in traits and environmental variables, we used Moran’s *I* correlograms (Legendre & Legendre, 2012) using the same custom-built spatial neighbourhood network.

### 2.6 Space-for-time substitution approach

We use this approach to make inference about microevolutionary processes. By using space as a substitute for time, we can use present environmental conditions to understand patterns produced by a past process (Wogan & Wang, 2018). Here, we sampled ponds along a rural-urban gradient (space) and calculated their rate of change in urbanisation from 1985 to 2021 (36 years) (time). Urbanisation rate varied from 0% to 2.11% per year. Ponds in the core urban had high urbanisation rates and had already undergone a moderate building process before 1985, providing evidence that ponds in urban core experienced a long lasting change. Thus, we assume frogs living in highly urbanised ponds have been experiencing the effect of urbanisation for more time than those in peri-urban or rural sites. Provided tropical treefrogs live approximately 12 years (Stark & Meiri, 2018), at last three generations have passed since 1985.

### 2.7 Data analysis

We checked for multicollinearity of all environmental variables using Variance Inflation Factor (VIF) in the R package usdm (Naimi et al., 2014), and standardized the data to zero mean and unit variance in the R package vegan (Oksanen et al., 2018). All variables had VIF < 6 and were retained for further analysis. Finally, the PC1 of Hill-Smith PCA (local environmental variables), MEM3 (spatial arrangement), urbanisation rate, and land surface temperature were entered into a Structural Equation Model to test for their direct and indirect effects on the mean and coefficient of variation of each phenotypic trait.

We calculate the mean as a sum of each phenotypic trait divided by the number of individuals in each pond. Also, we calculated the Kvålseth coefficient of variation (KCV; see Kvålseth, 2017) for each phenotypic trait as the standard deviation divided by the square root of the mean of squared values, with an R function provided by Lobry et al., (2023). Path analysis was performed in the R package lavaan (Rosseel, 2012). Figures were made in the R package semPlot (Epskamp, 2022).

To estimate the adaptive landscape, we separately regressed the Scaled Mass Index as a function of body size, size-free leg length, and head shape. We used a non-parametric regression technique called projection-pursuit regression implemented in the gsg R package (Morrissey, 2014). This approach assumes that the partial regression coefficient represents the selection gradient as a way to estimate the adaptive landscape (Lande & Arnold, 1983). The data and associated R code are available in Dryad (Severgnini & Provete 2024).

## 3. RESULTS

We sampled 768 individuals of *Dendropsophus nanus* along an urban gradient (∼ 30 individuals per pond). Urbanisation rate and the spatial structure were directly, positively related to local heterogeneity in all models (Table 1). Mean body size was positively related to PC1, which means it increased in ponds with excavated margins and decreased in those with temporary hydroperiod, invertebrate predators, and cattle (Table 1; Figure 3; Figure S7, S8, S9). Mean body size was negatively associated with MEM3, which means it decreased from the periphery to the urban centre (contrast Figure S6 with Figure S8). Mean head shape was negatively associated with local environmental heterogeneity and positively related to MEM3. This means that frogs in Northeast-Southwest distributed ponds had short and narrow heads, while frogs with long and wide heads were found in ponds with excavated banks. Body condition was partially, negatively associated with environmental heterogeneity, meaning that frogs with lower body condition occurred in ponds with excavated margins, whereas temporary ponds with invertebrate predators and cattle had frogs with better body condition (Table 1; Figure 3; Figure S7, S8).

**Figure 3.**
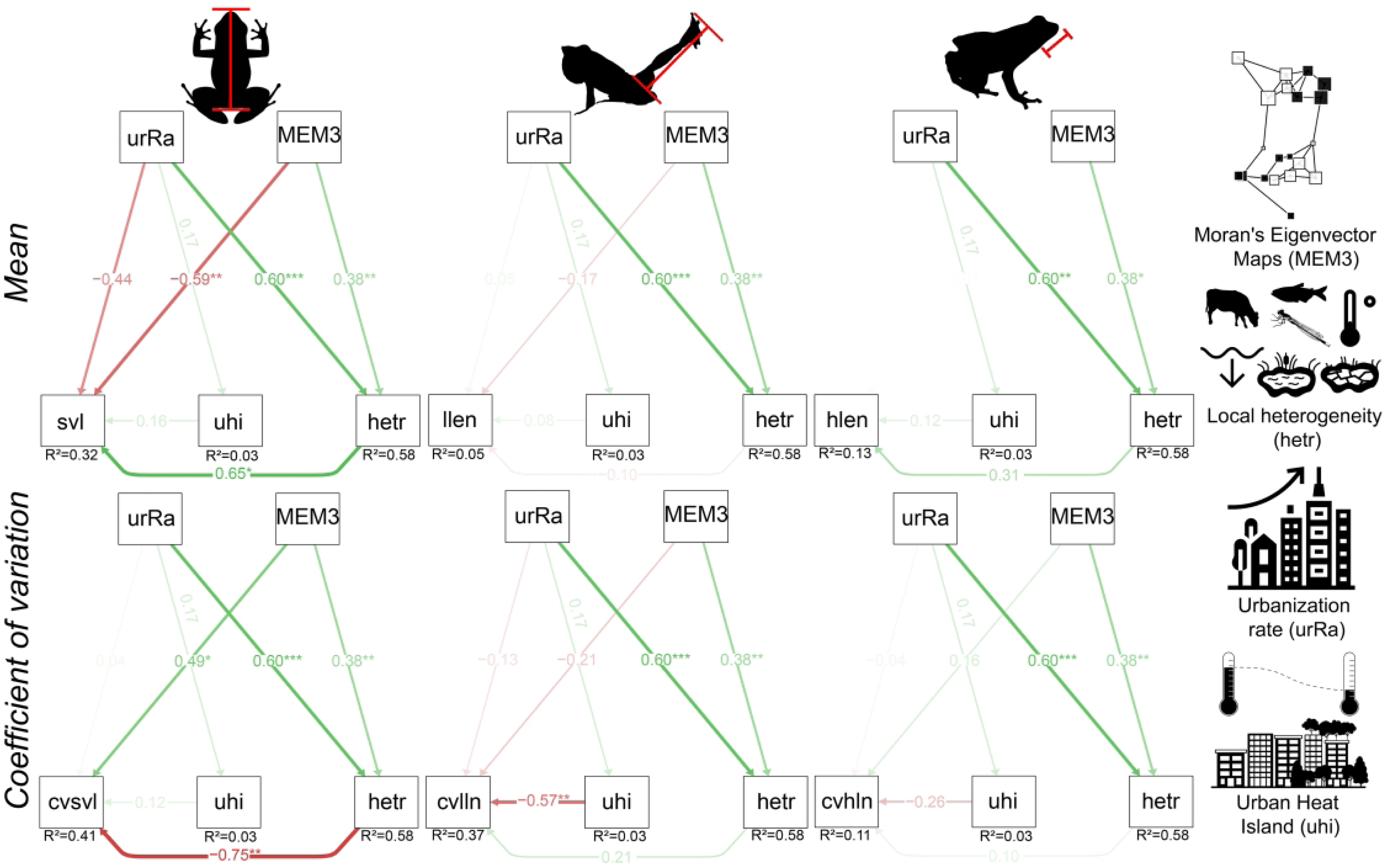
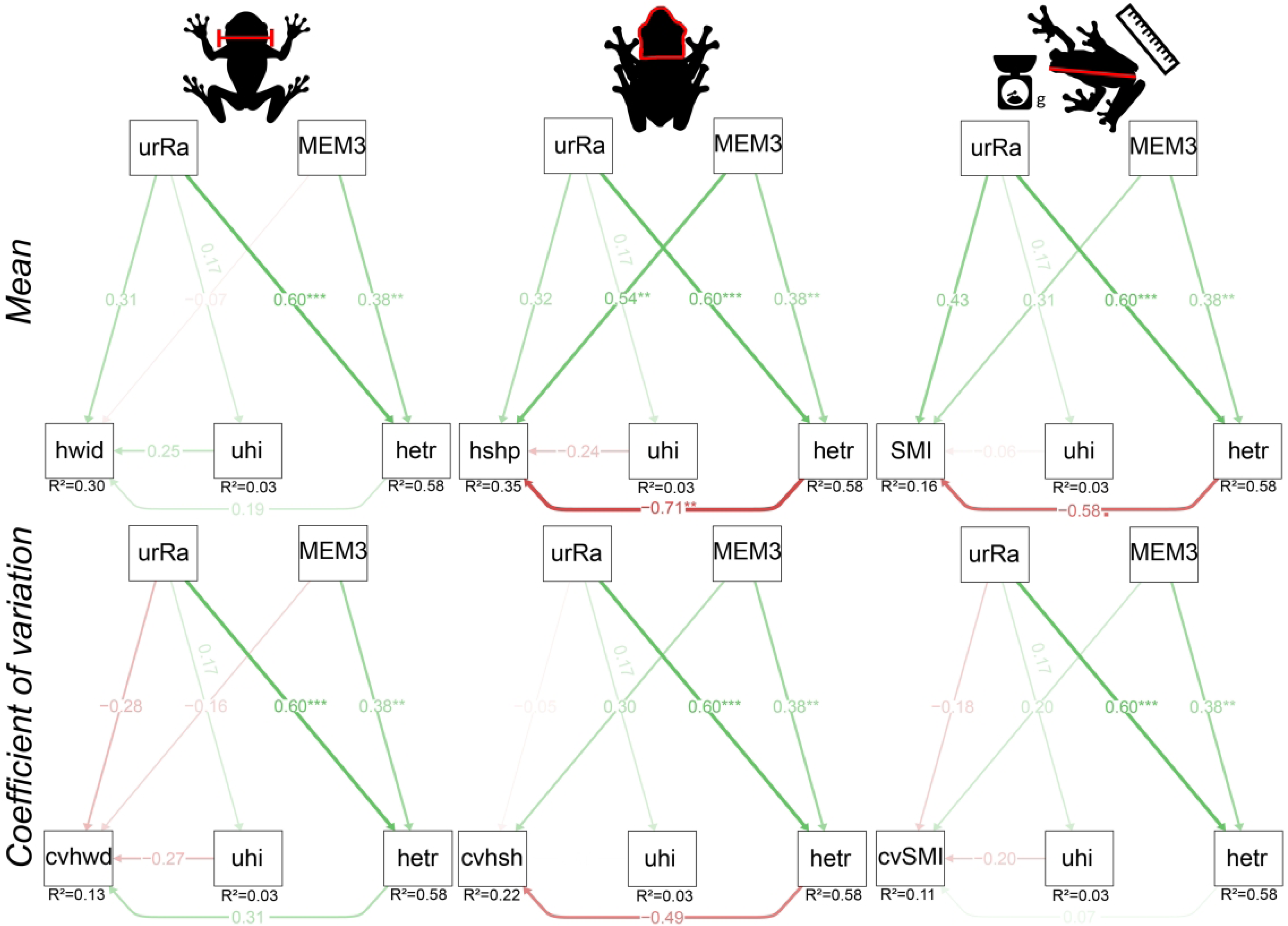
Path diagrams showing the standardized coefficients (β) of models testing the influence of predictor variables: Urbanization rate (urRa); Moran eigenvector maps (MEM3); Land Surface Temperature – Urban heat island (uhi), and Local heterogeneity (hetr) – (first axis of a Hill-Smith Principal Component Analysis) on mean and coefficient of variation of body size (svl); leg length (llen); and head length (hlen). Goodness fit: χ^2^= 1.115; DF=2; P = 0.573. Frogs, cow, fish, and dragonfly silhouettes are CC-BY from PhyloPic. Local heterogeneity, urbanization rate, and urban heat island silhouettes are CC-BY 3.0 from Noun Project (ponds created by Ircham; thermometer created by Andi Nur Abdillah, pond depth created by Alessandro Suraci, urban heat island created by Softscape, and urbanization rate created by WiStudio).

**Table 1.** Results of Structural Equation Models showing models results about the influence of predictor variables: Local heterogeneity (hetr) – (first axis of Hill-Smith Principal Component Analysis); Urbanisation rate (urRa); Urban heat island – Land Surface Temperature (uhi); and Moran’s Eigenvector Maps (MEM3) on mean body size; mean leg length; mean head width; mean head length; mean head shape; and Scaled Mass Index. We also tested the effects of the same predictors on the Kvålseth coefficient of variation (KCV) of phenotypic traits. The last subheading shows paths present in all models with the same results. Goodness of fit: χ^2^ = 1.115, DF = 2, *P* = 0.573. Std. β = Standardized regression coefficients; z-value = z statistics; *P* (>|z|) = probability value; Est. (β) = regression coefficients; Std. Err = Standard error. Significant *P* values are in bold.

| | Est. ($\beta$) | Std. Err | z-value | $P(>\vert z\vert )$ | Std. $\beta$ |
| --- | --- | --- | --- | --- | --- |
| *Mean* |  |  |  |  |  |
| **Body size** | ~ |  |  |  |  |
| Local heterogeneity | 0.300 | 0.127 | **2.363** | **0.018** | **0.658** |
| Urbanisation rate | -0.205 | 0.114 | -1.803 | 0.071 | -0.449 |
| Urban heat Island | 0.076 | 0.083 | 0.911 | 0.362 | 0.166 |
| MEM3 | -0.269 | 0.096 | **-2.792** | **0.005** | **-0.589** |
| **Leg length** | ~ |  |  |  |  |
| Local heterogeneity | -0.027 | 0.096 | -0.278 | 0.781 | -0.091 |
| Urbanisation rate | 0.017 | 0.086 | 0.196 | 0.844 | 0.058 |
| Urban heat Island | 0.026 | 0.063 | 0.408 | 0.683 | 0.088 |
| MEM3 | -0.052 | 0.073 | -0.709 | 0.478 | -0.177 |
| **Head width** | ~ |  |  |  |  |
| Local heterogeneity | 0.014 | 0.020 | 0.680 | 0.496 | 0.192 |
| Urbanisation rate | 0.022 | 0.018 | 1.247 | 0.213 | 0.315 |
| uhi | 0.018 | 0.013 | 1.358 | 0.175 | 0.251 |
| MEM3 | -0.005 | 0.015 | -0.323 | 0.747 | -0.069 |
| **Head length** | ~ |  |  |  |  |
| Local heterogeneity | 0.023 | 0.022 | 1.006 | 0.314 | 0.317 |
| Urbanisation rate | 0.00 | 0.020 | 0.015 | 0.988 | 0.004 |
| Urban heat Island | 0.009 | 0.015 | 0.601 | 0.548 | 0.124 |
| MEM3 | 0.001 | 0.017 | 0.065 | 0.948 | 0.016 |
| **Head shape** | ~ |  |  |  |  |
| Local heterogeneity | -0.015 | 0.006 | **-2.626** | **0.009** | **-0.714** |
| Urbanisation rate | 0.007 | 0.005 | 1.312 | 0.19 | 0.319 |
| Urban heat Island | -0.005 | 0.004 | -1.387 | 0.165 | -0.247 |
| MEM3 | 0.011 | 0.004 | **2.646** | **0.008** | **0.545** |
| **Scaled mass index** | ~ |  |  |  |  |
| Local heterogeneity | -0.027 | 0.014 | **-1.892** | **0.058** | **-0.587** |
| Urbanisation rate | 0.02 | 0.013 | 1.569 | 0.117 | 0.435 |
| Urban heat Island | -0.003 | 0.009 | -0.320 | 0.749 | -0.065 |
| MEM3 | 0.014 | 0.011 | 1.322 | 0.186 | 0.311 |
| *Coefficient of variation* |  |  |  |  |  |
| **cv Body size** | ~ |  |  |  |  |
| Local heterogeneity | -0.007 | 0.003 | **-2.89** | **0.004** | **-0.751** |
| Urbanisation rate | 0.00 | 0.002 | 0.194 | 0.846 | 0.045 |
| Urban heat Island | 0.001 | 0.002 | 0.720 | 0.471 | 0.122 |
| MEM3 | 0.005 | 0.002 | **2.486** | **0.013** | **0.489** |
| **cv Leg length** | ~ |  |  |  |  |
| Local heterogeneity | 0.003 | 0.004 | 0.778 | 0.437 | 0.209 |
| Urbanisation rate | -0.002 | 0.003 | -0.523 | 0.601 | -0.126 |
| Urban heat Island | -0.008 | 0.002 | **-3.263** | **0.001** | **-0.574** |
| MEM3 | -0.003 | 0.003 | -1.031 | 0.303 | -0.210 |
| **cv Head width** | ~ |  |  |  |  |
| Local heterogeneity | 0.006 | 0.006 | 0.988 | 0.323 | 0.311 |
| Urbanisation rate | -0.005 | 0.006 | -0.986 | 0.324 | -0.278 |
| Urban heat Island | -0.005 | 0.004 | -1.324 | 0.185 | -0.273 |
| MEM3 | -0.003 | 0.005 | -0.704 | 0.482 | -0.168 |
| **cv Head length** | ~ |  |  |  |  |
| Local heterogeneity | 0.002 | 0.005 | 0.314 | 0.753 | 0.101 |
| Urbanisation rate | -0.001 | 0.005 | -0.164 | 0.870 | -0.047 |
| Urban heat Island | -0.004 | 0.003 | -1.227 | 0.220 | -0.257 |
| MEM3 | 0.003 | 0.004 | 0.658 | 0.510 | 0.160 |
| **cv Head shape** | ~ |  |  |  |  |
| Local heterogeneity | -0.111 | 0.067 | -1.651 | 0.099 | -0.493 |
| Urbanisation rate | -0.012 | 0.06 | -0.192 | 0.848 | -0.051 |
| Urban heat Island | 0.001 | 0.044 | 0.013 | 0.990 | 0.003 |
| MEM3 | 0.068 | 0.051 | 1.330 | 0.184 | 0.301 |
| **cv Scaled mass index** | ~ |  |  |  |  |
| Local heterogeneity | 0.005 | 0.020 | 0.245 | 0.807 | 0.078 |
| Urbanisation rate | -0.012 | 0.018 | -0.661 | 0.509 | -0.188 |
| Urban heat Island | -0.013 | 0.013 | -0.961 | 0.337 | -0.200 |
| MEM3 | 0.013 | 0.015 | 0.843 | 0.399 | 0.204 |
| *Predictor variables* |  |  |  |  |  |
| **Urban Heat Island** | ~ |  |  |  |  |
| Urbanisation rate | 0.172 | 0.215 | 0.800 | 0.423 | 0.172 |
| **Local heterogeneity** | ~ |  |  |  |  |
| Urbanisation rate | 0.600 | 0.143 | **4.202** | **0.000** | **0.600** |
| MEM3 | 0.382 | 0.143 | **2.676** | **0.007** | **0.382** |

The coefficient of variation (CV) of body size was negatively related to environmental heterogeneity and positively with MEM3. This means that the body size of frogs varied less in ponds with excavated margins, whereas it varied more in temporary ponds with invertebrate predators and cattle. Ponds in Northeast-Southwest regions had higher variation in body size than those in the Northwest-Southeast region (Table 1; Figure 3; Figure S7, S8). The CV of leg length was negatively affected by land surface temperature (Table 1; Figure 3; Figure S7, S8). The remaining traits were not directly affected by any predictor variable. Finally, neither traits nor predictor variables were spatially autocorrelated (Figure S10 and S11). Interestingly, we found no direct effect of urbanisation rate on any trait, except for a small negative effect on mean body size and a positive one on mean body condition. Instead, its effect on the mean and variance body size and mean head shape seems to be indirect, via pond environmental variables.

Body size showed a negative, though weak, directional selection (β = - 0.12; *P* < 0.001), while head shape was under weak, positive directional selection (β = 2.89; *P* < 0.002), in which individuals with wider and longer heads had increased fitness (Figure S12). The linear selection coefficient was not significant for leg length. Likewise, the quadratic coefficient was not significant for any trait (Table S2).

## 4. DISCUSSION

Our results showed that mean body size, head shape, and the variation in body size were directly influenced, but in opposite ways, by the spatial arrangement of ponds and local environmental heterogeneity. The variation in leg length decreased with increasing land surface temperature. Lastly, fitness decreased with increasing body size, while it increased in frogs with wider and longer heads. Contrarily to our initial hypotheses, we did not find a direct effect of urbanisation rate on phenotypic traits, except for a small effect on mean body size and body condition. Instead, urbanisation influenced phenotypes indirectly via local environmental variables.

Mean body size increased in ponds with excavated margins and decreased in those with temporary hydroperiod, cattle and vertebrate predators. Urbanisation rate was also positively related to local heterogeneity. These results suggest that urbanisation is indirectly influencing mean body size via changes in pond environmental characteristics. *Dendropsophus nanus* is a small treefrog that usually calls on tree branches or leaves on pond margins (Menin et al., 2005). Man-made ponds in highly urbanized areas might have less appropriate microhabitats (Hutto & Barrett, 2021) and calling site availability (Hamer & McDonnell, 2008). Thus, frog populations may be in turn responding to these environmental filters by altering their phenotypic traits. Most previous studies found smaller body sizes in urban environments compared to rural or undisturbed sites (e.g., Jennette et al., 2019; Komine et al., 2022; Liu et al., 2021; Matías-Ferrer & Escalante, 2015). Also, Liu et al., (2021) found that larger species were more tolerant to anthropogenic habitat modification, but this pattern was reversed after controlling for phylogenetic relationships. The fact that the mean body size of frogs is responding more strongly to environmental changes driven by urbanisation, instead of temperature-related variables like urban heat island suggests that physiological constraints seem less important than phenotype-habitat matching (Edelaar et al. 2008) in young cities. This is somewhat unexpected, since frogs have a narrow thermal performance (see Wells 2007; Hillman et al., 2008). However, the relatively recent urbanisation process might not yet be strong enough to induce local adaptations.

In contrast, the variation in body size was negatively related to environmental heterogeneity, decreasing in man-made ponds. Ponds in Northeast-Southwest regions (ie. most of the urbanised ponds) tend to have frogs with more variable body sizes, but this spatial pattern was less strong. This pattern is similar to what previous studies found for mean body size (e.g., Jennette et al., 2019; Komine et al., 2022). Artificial ponds might be more simplified environments (Hutto & Barrett, 2021), which can constrain the variance of body size around an optimal value. This result also suggests that the phenotype of this frog population might be experiencing ongoing, yet weak, natural selection. Thus, body size seems to have idiosyncratic responses to urbanisation (Langerhans & Kern, 2020) and differ between tropical and temperate cities. Our study adds new evidence indicating reduced phenotypic variability in urban areas.

Treefrogs had narrow and short heads in mean in man-made ponds. This phenotypic response might be due to the reduced size and low prey diversity in urban ponds (Santana et al., 2019), which would elicit an adaptive response in frogs to decrease their heads to become more efficient in consuming small prey. Mouth width and head size are the main traits influencing the size (Parmelee, 1999), amount, and diversity of prey consumed by frogs (Moroti et al., 2021). Therefore, narrower heads in frog populations might be a phenotypic response to the decrease in body size of their prey in urban sites (e.g., Ishitani et al., 2003; Merckx et al., 2018; Piano et al., 2020; Ulrich et al., 2008). Therefore, mean head shape showed a different pattern compared to body size, increasing from Northeast-Southwest to Northwest-Southeast, suggesting that different selective pressures at fine spatial scales (e.g., pond level) can be acting on each phenotypic trait.

The CV of leg length decreased with increasing land surface temperature. Environmental stressing factors, such as increased surface temperature, can promote developmental disruptions (Thompson et al. 2022) that reduce phenotypic variance. A meta-analysis found that temperature variation in the larval stage had a small, yet significant effect on the leg length of adult frogs due to disruptions in developmental, but not growth rate (Tejedo et al. 2010). Differences in the timing of developmental events, or heterochrony, can produce significant differences in relative leg length (Emerson 1986). Increasing temperature usually decreases relative humidity, which poses a further challenge to frogs in dealing with evaporative water loss. Frogs can avoid water loss by adopting a water-conserving posture during diurnal sleeping (Mitchell & Bergmann, 2016), in which they hide their limbs against their body to reduce their surface area. Habitat selection and behaviours to avoid dissection seem not to affect their jumping performance (Mitchell & Bergmann, 2016). Therefore, lower variance of leg length in ponds with higher temperatures are likely due to disruptions in developmental rate (Tejedo et al. 2010) and can constrain the evolution of this phenotype, with potential consequences for thermoregulatory physiology and performance.

Neither trait nor environmental variable exhibited a clear spatial autocorrelation pattern. These results suggest that we analysed a pure environmental gradient and the trait-environment relationships found are not confused with spatial autocorrelation (Urban, 2011). Furthermore, it provides evidence that there is no local adaptation in any phenotypic trait, suggesting that populations still maintain a moderate-to-high gene flow (see Urban, 2011). Some amphibian populations can maintain gene flow in urbanized areas, preserving genetic diversity (e.g., Fusco et al., 2020). Consequently, moderate gene flow can prevent local adaptation (Miles et al., 2019; Rivkin et al., 2019). Our study tested how phenotypic traits differentiate along rural–urban sites while accounting for their environmental characteristics and spatial arrangement to make evolutionary inferences. The absence of a clear spatial autocorrelation in environmental variables also allows separating adaptive from neutral responses of phenotypes, akin of the sampling design used in Gilbert & Lechowicz (2004) for testing neutral versus niche-based processes in plant communities. Thus, our sampling design allowed us to disentangle the isolated effects of space and environment on the variation of phenotypic traits.

Both body size and head shape were under weak directional selection, but in opposite directions. Also, scaled mass index was only partially, negatively affected by local environmental heterogeneity. Our results suggest that larger frogs have lower fitness. Consequently, natural selection seems to disfavour an increase in body size in more urbanized ponds. This result is counterintuitive, because larger animals should have higher fitness, since larger body sizes translates into higher competitive ability, dispersal, food acquisition, and consequently higher fecundity and survivorship (Blanckenhorn, 2000; Smith & Belk, 2018). This result suggest that frogs might be experiencing maladaptation in terms of their body size (Brady et al., 2019a; 2019b), likely due to urban heat island (Diamond & Martin, 2020a, 2020b). As discussed above, populations can still be experiencing some degree of gene flow, which might promote trait homogenization, producing less fit phenotypes (Urban, 2011). However, phenotypic traits can evolve even when fitness rewards are low (Fisher & McAdam, 2019). The variation in trait values in a population, with some individual having higher trait values that improve their fitness, while others have lower trait values, may render individuals more or less adapted to novel environmental conditions (Hendry, 2017) found in urban sites. Another interesting result is that individuals with broader and longer heads had higher fitness, likely because they allow the acquisition of a wider range of resources (Emerson, 1985b).

## 5. CONCLUSION

Most results did not agree with our hypotheses, except for CV of body size that decreased with urbanisation, CV of leg length that decreased with land surface temperature. Our results suggest that the lack of a clear spatial variation in phenotypic traits can be due to weak selection as a result of a recent, although intense, urbanisation process. As such, populations might be experiencing moderate-to-high gene flow that prevents local adaptation. In conclusion, eco-evolutionary dynamics in tropical cities seem to have a different pace compared to temperate ones, producing less significant phenotypic changes between rural and urban frog populations. Our results can contribute to building urban ecological theory that explicitly includes city age, their development, growth rate, and history.

## Author Contributions

MRS: Writing–Original Draft (Leading), Methodology (Equal), Data Curation, Formal Analysis.

DBP: Conceptualization (Leading), Methodology (Equal), Writing–Reviewing and Editing (Equal), Supervision. All authors contributed critically to the drafts and gave final approval for publication.

## Acknowledgements

Mauricio Vancine kindly adapted an R code used to extract buffers from raster automatically. We thank landowners for allowing access to their properties. Heloísa M. Rodrigues, Marciane R. Severgnini, Bruna C. Yoshida, Philip T. Soares, Adriana C. Acero-Murcia, Nicolle Prado, Klysman Fernandes, Bruno Fines, Letícia A. da Cruz, Eduardo Morel, and Ana Torres helped with field work. Megan J. Thompson kindly provided feedback on an earlier version of the manuscript. We dedicate this study to Prof. Marcelo Menin, who passed away due to COVID-19 in 2021.

## Conflict of interest

The authors declare no conflict of interest.

## Data availability

All data and associated R code used to run the analysis of this manuscript are available for peer review at DataDryad https://datadryad.org/stash/share/Ev401eLzTpaPwnevzQGPAU7lwO8jZiUMlfqk rjRXYpA.

## Funding

DBP is supported by a grant from the Brazilian National Council on Research and Technological development – CNPq (#407318/2021-6). This study was funded in part by the Coordenação de Aperfeiçoamento de Pessoal de Nível Superior – Brasil (CAPES) – Finance Code 001 to MRS and DBP. DBP received a Humboldt Foundation fellowship for experienced researchers during the final stages of writing.

## Ethics approval

ICMBio provided collecting permits (#80075-1). This study was approved by the ethics committee of our university (#1.203/2021).

## Supplementary material for

**Table S1.** Local environmental variables used to compute Hill-Smith Principal Component Analysis and their loadings along the first two axes. The first axis was used to represent local heterogeneity.

| Variables | CS1 | CS2 |
| --- | --- | --- |
| Hydroperiod permanent | 0.146 | 0.058 |
| Hydroperiod temporary | -0.878 | -0.347 |
| Margin profile excavated | 0.634 | -0.217 |
| Margin profile flat | -0.330 | 0.613 |
| Margin profile sloping | -0.304 | -0.396 |
| Predator both (vertebrate and invertebrate) | 0.298 | -0.065 |
| Predator invertebrate | -0.596 | 0.131 |
| Cattle absent | 0.196 | 0.165 |
| Cattle present | -0.628 | -0.528 |
| Floating vegetation (%) | -0.341 | 0.381 |
| Temperature (°C) | -0.319 | -0.431 |
| Area (m<sup>2</sup>) | 0.012 | 0.577 |
| Pond depth (mean, cm) | 0.380 | -0.160 |

**Table S2.** Results of the adaptive landscape for size-free leg length, body size, and head shape showing linear selection coefficients (β) and quadratic coefficients (γ). SE – Standard error; Estimates – Slope. Significant *P* values are in bold.

| Trait | Selection coefficient | estimates | SE | $P$ value |
| --- | --- | --- | --- | --- |
| Leg length | Beta | 0.04693604 | 0.04943377 | 0.348 |
|  | Gamma | 0.00000015 | 0.00000061 | 0.376 |
| Body size | Beta | -0.1201279 | 0,01997591 | **0.001** |
|  | Gamma | 0.00000000 | 0.00000066 | 0.382 |
| Head shape | Beta | 2.892416 | 0.811965 | **0.002** |
|  | Gamma | -0.00000007 | 0.00000082 | 0.972 |

**Figure S1.**
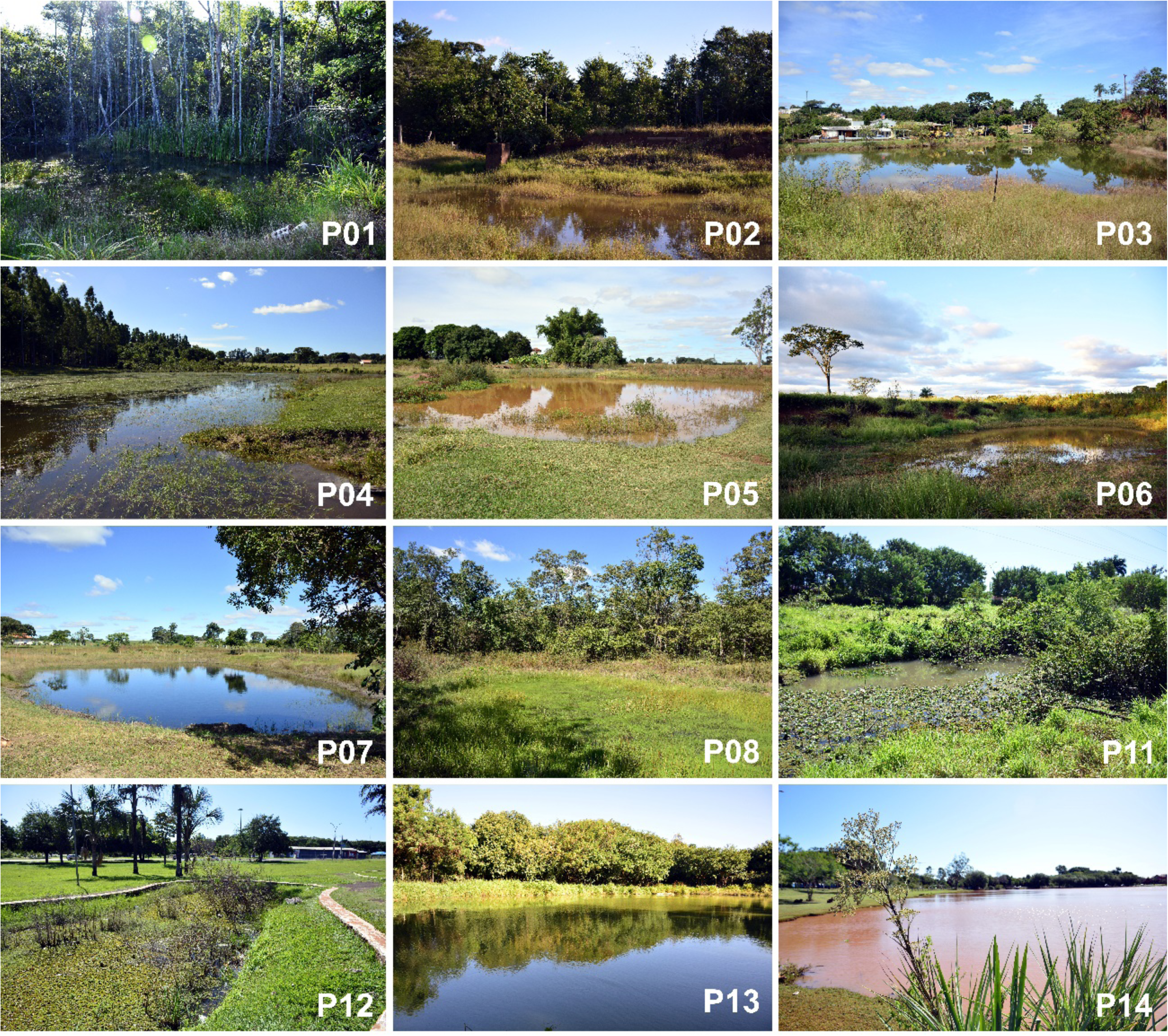
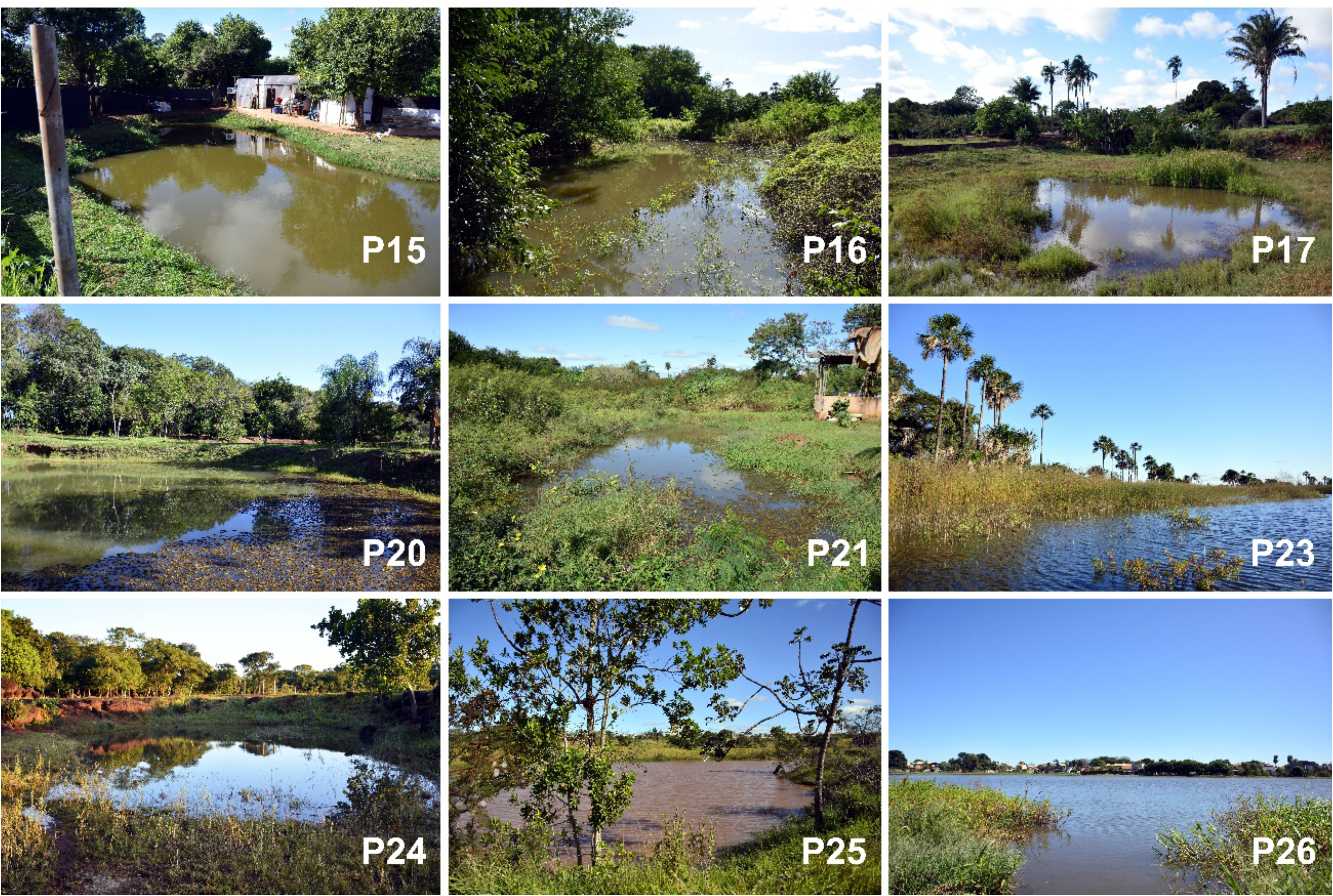
Ponds sampled along the rural–urban gradient in Campo Grande, Mato Grosso do Sul, Brazil. P11 and P21 are examples of ponds with excavated margins, P23 had a flat margin, while P04 and P05 had slopping margins.

**Figure S2.**
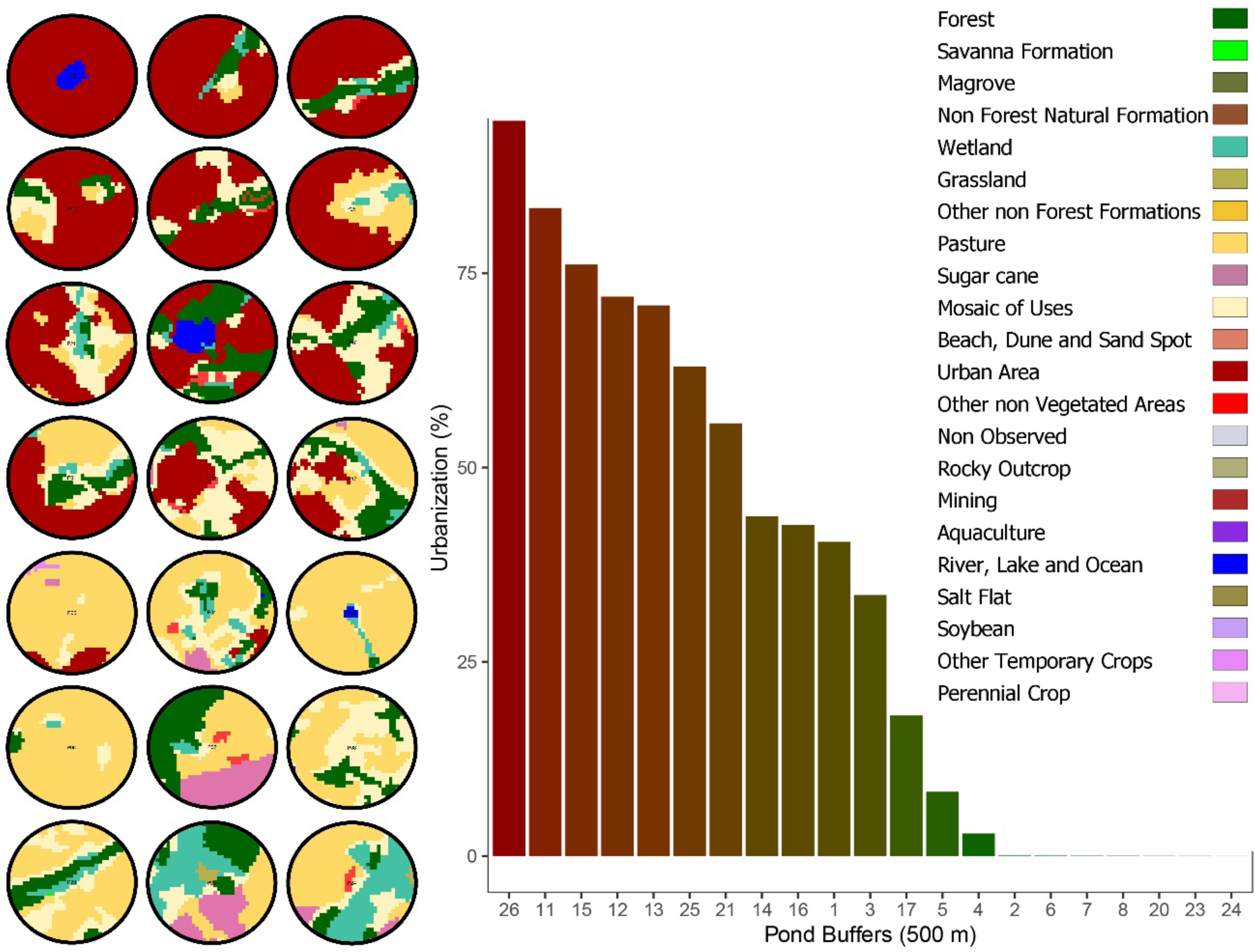
Percentage of impervious surface (roads and buildings) as of 2021within a 500-m buffer around the 21 sampled ponds along the urbanization gradient.

**Figure S3.**
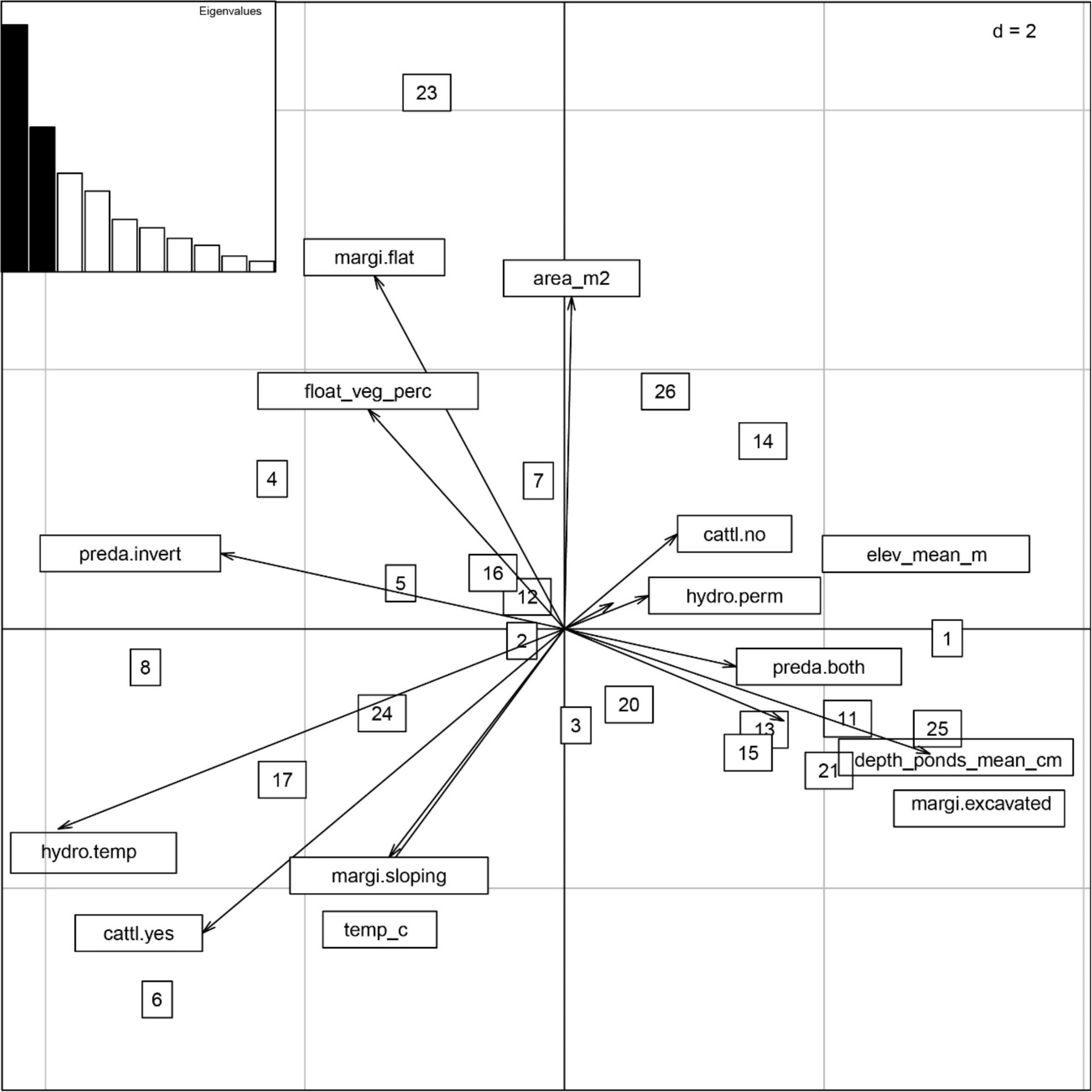
Ordination diagram of the first two axes of the Hill-Smith Principal Component Analysis showing the ponds (numbers in squares) and local environmental variables (arrows).

**Figure S4.**
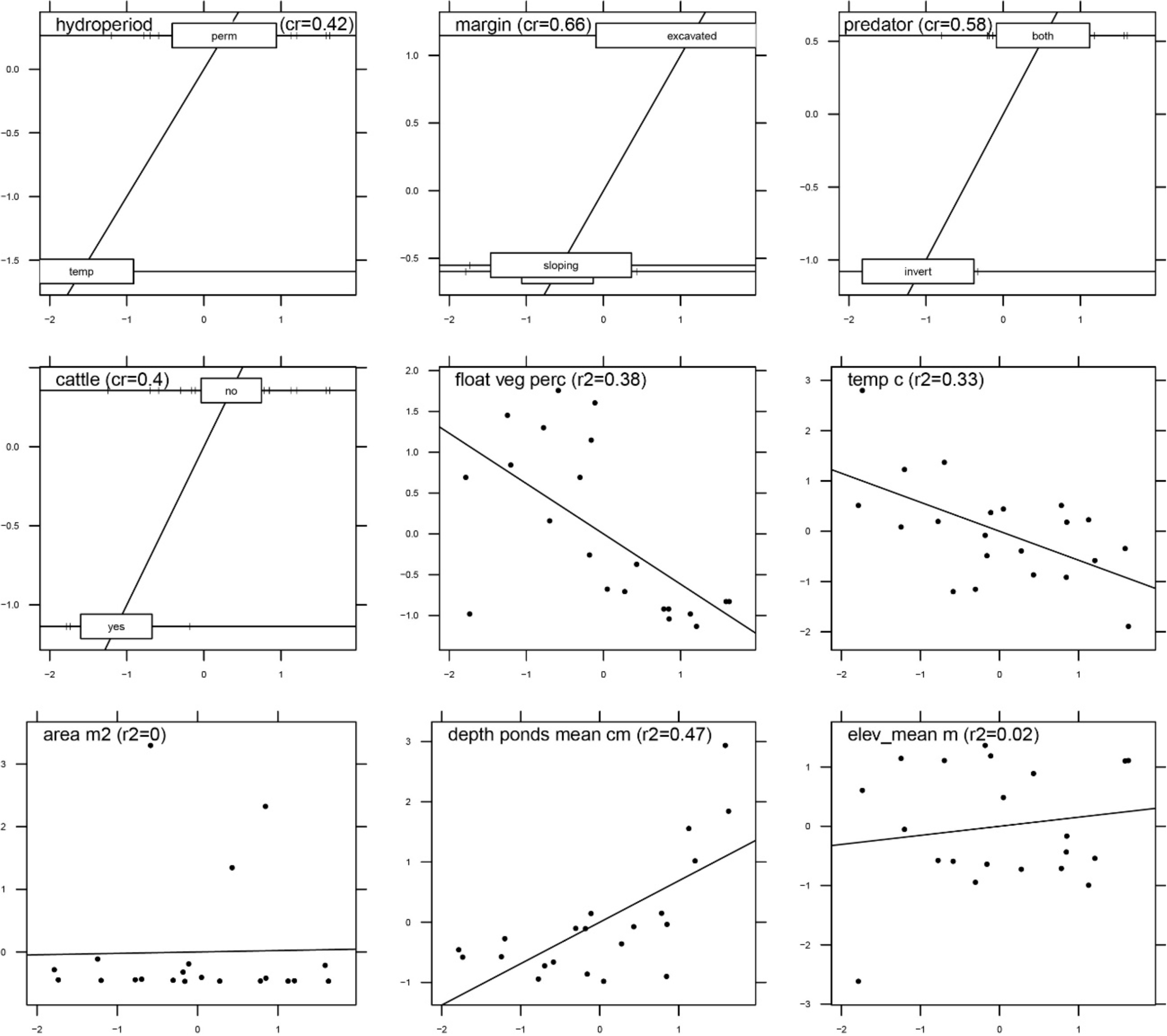
Scores of the local environmental variables along the first axis of the Hill-Smith Principal Component Analysis. For the quantitative variables r^2^ = squared regression coefficients, and for discrete variables cr = correlation ratio.

**Figure S5.**
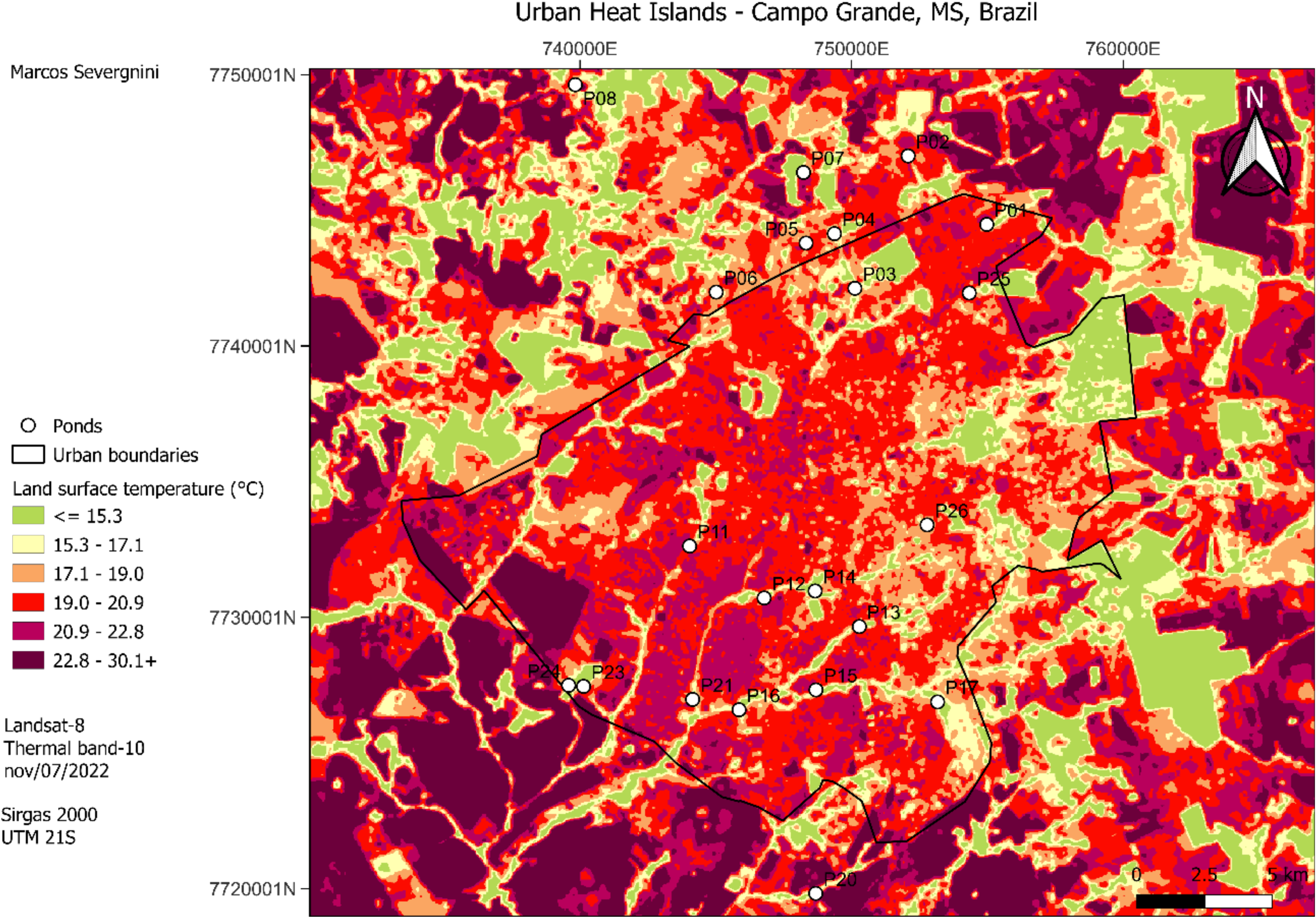
Map of land surface temperature/urban heat islands of Campo Grande, Mato Grosso do Sul, Brazil. White circles represent sampling sites (ponds). Map features LANDSAT 8 thermal band 10 extracted from United States Geological Survey (USGS). Black line delimits the urban perimeter extracted from https://sisgran.campogrande.ms.gov.br/; and prepared on QGIS v. 3.22.1.

**Figure S6.**
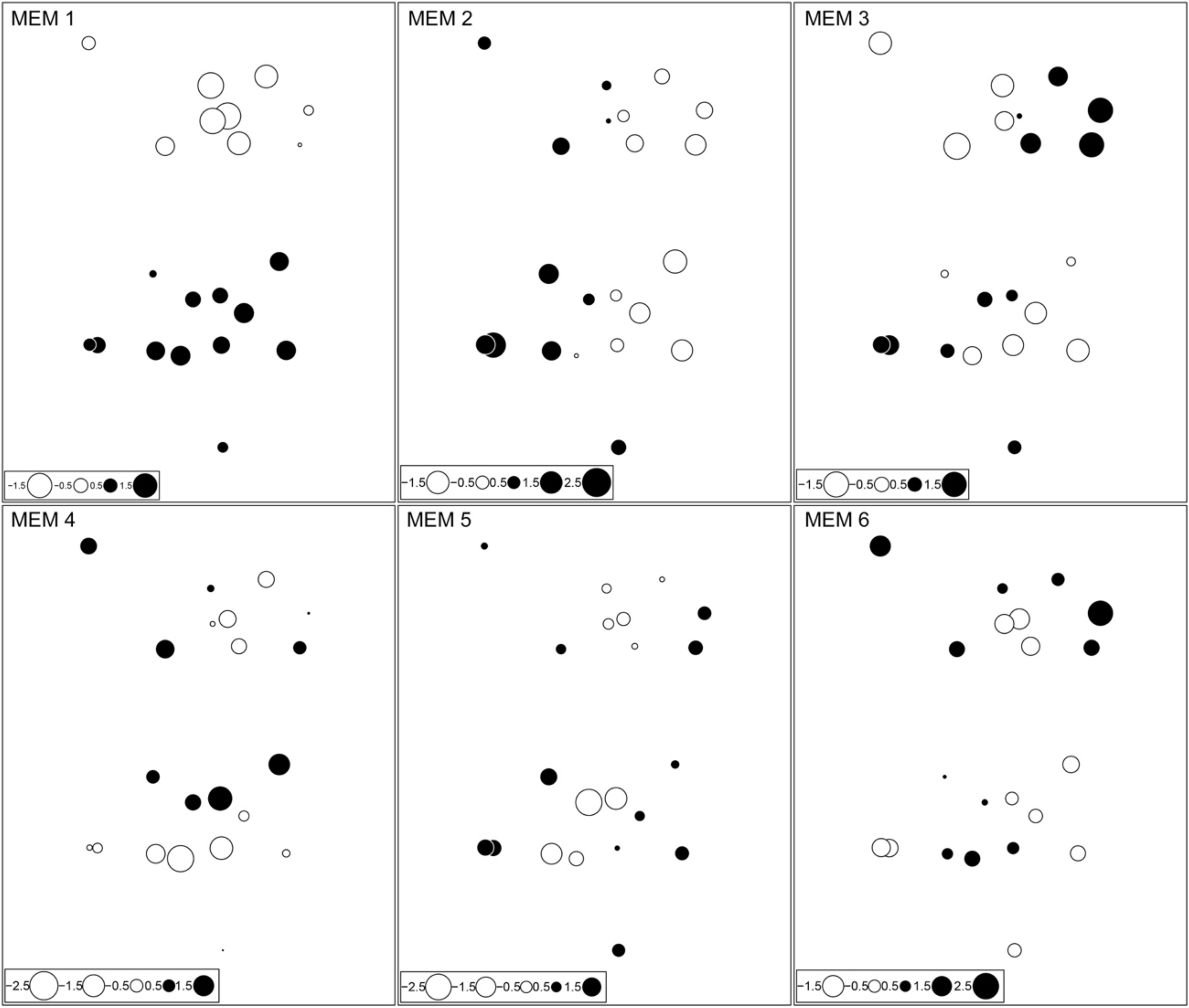
The six Moran Eigenvector Maps (MEMs) generated with the custom spatial neighbourhood network that have positive (Moran’s *I* > 0) and significant spatial autocorrelation. Each dot represents a sampled pond. Black dots represent positive and white dots represent negative scores along each eigenvector. Each eigenvector describes the spatial arrangement of ponds in a given spatial scale.

**Figure S7.**
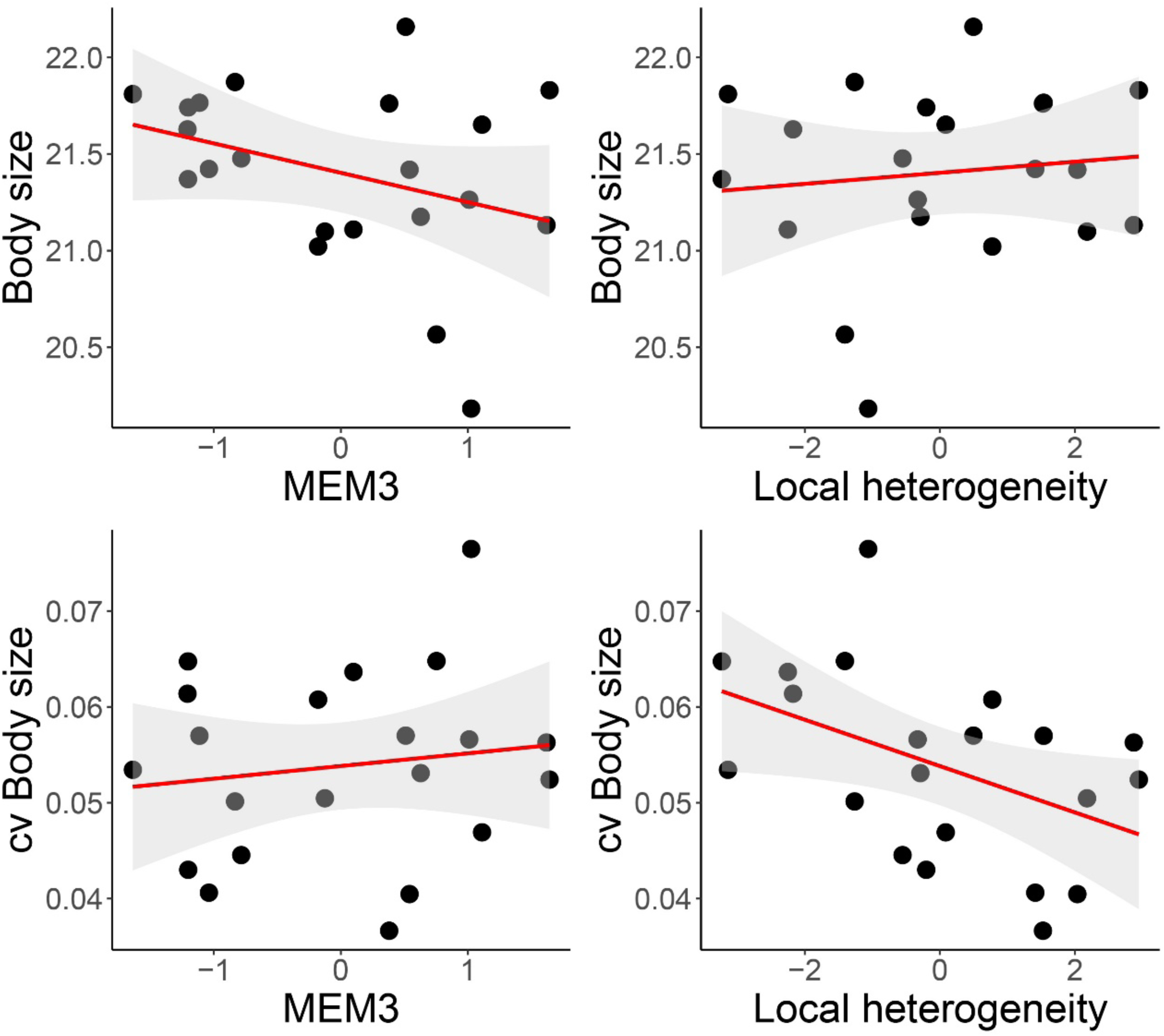
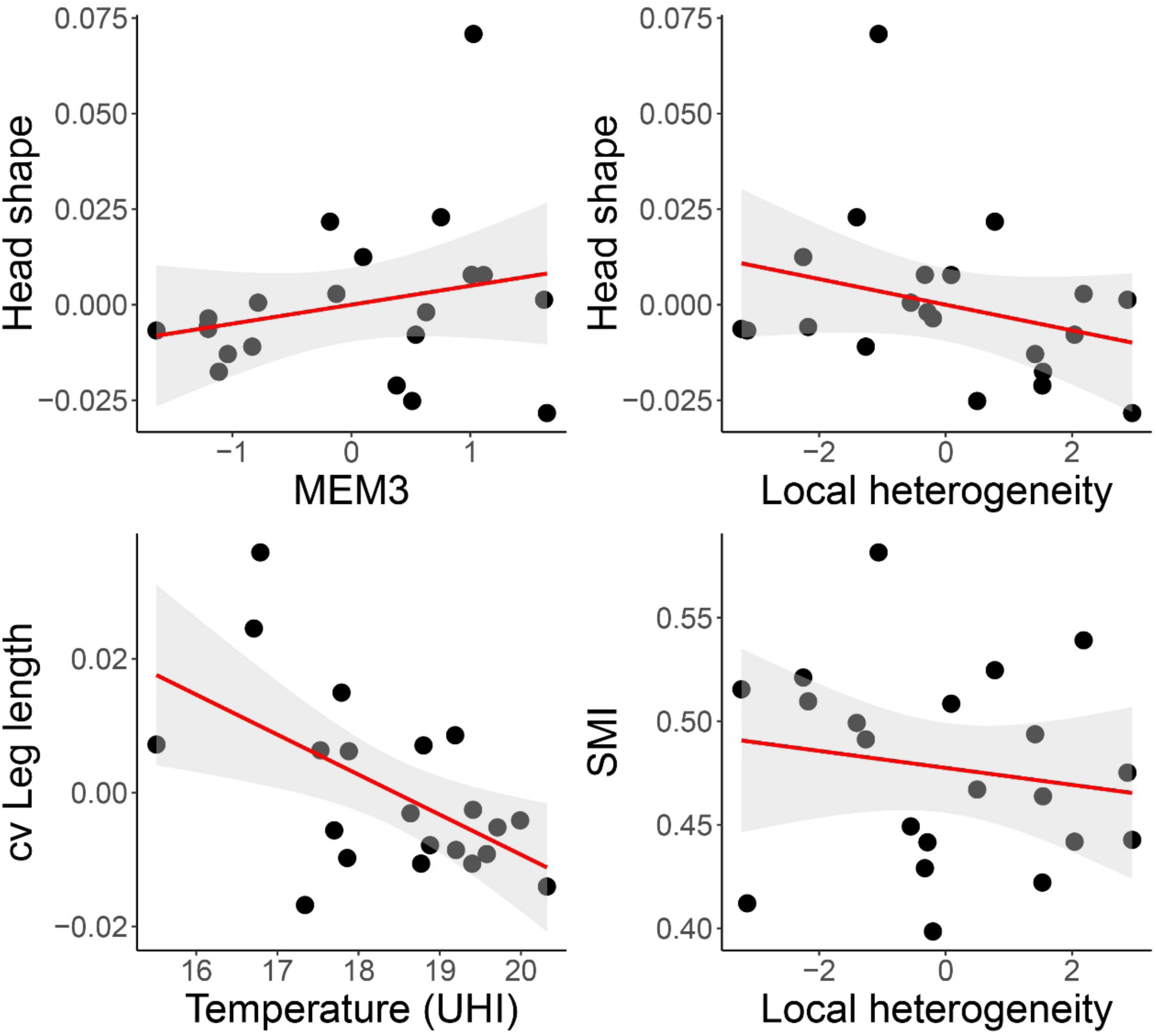
Scatter plots showing the relationship between variables that were significant in the structural equation model. MEM3: Moran Eigenvector Maps (MEMs); cv: coefficient of variation; UHI: urban heat island – land surface temperature (next panel).

**Figure S8.**
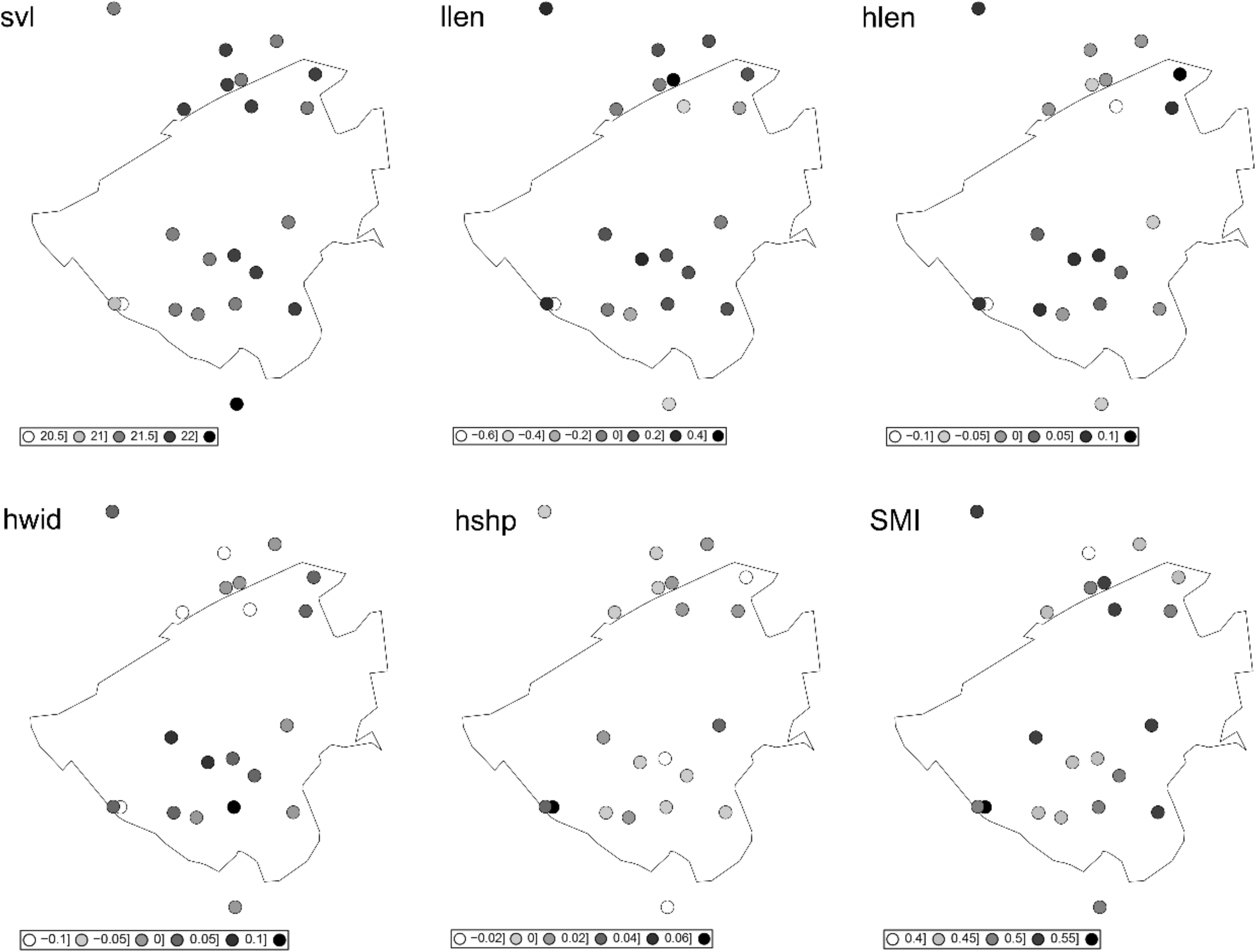
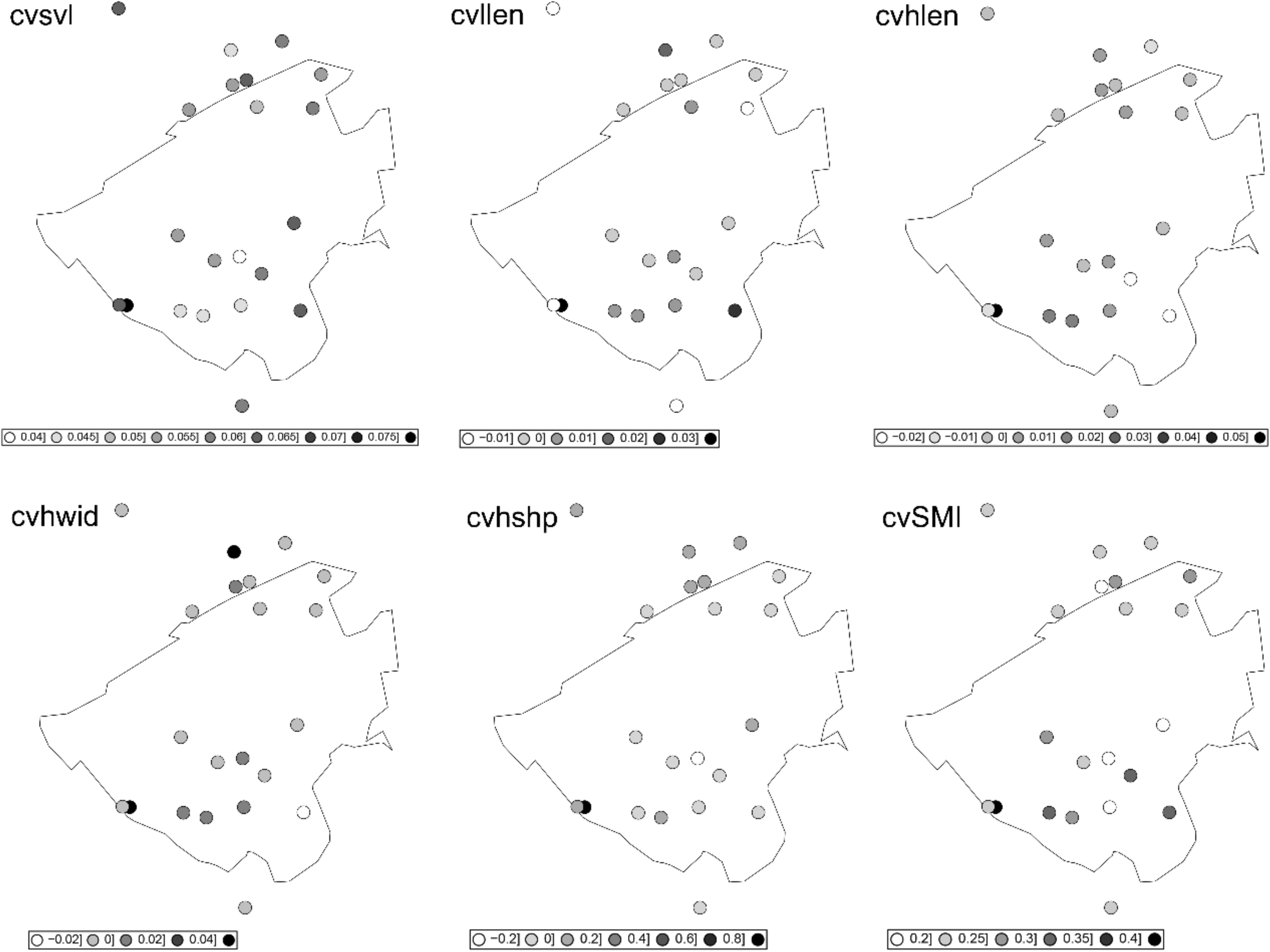
Mean and coefficient of variation of body size (svl); leg length (llen); head length (hlen); head width (hwid); head shape (hshp); and Scaled mass index (SMI) across urban gradient of Campo Grande city. Circles represents trait values that are presented in grey gradient ranging from min to max values of each trait.

**Figure S9.**
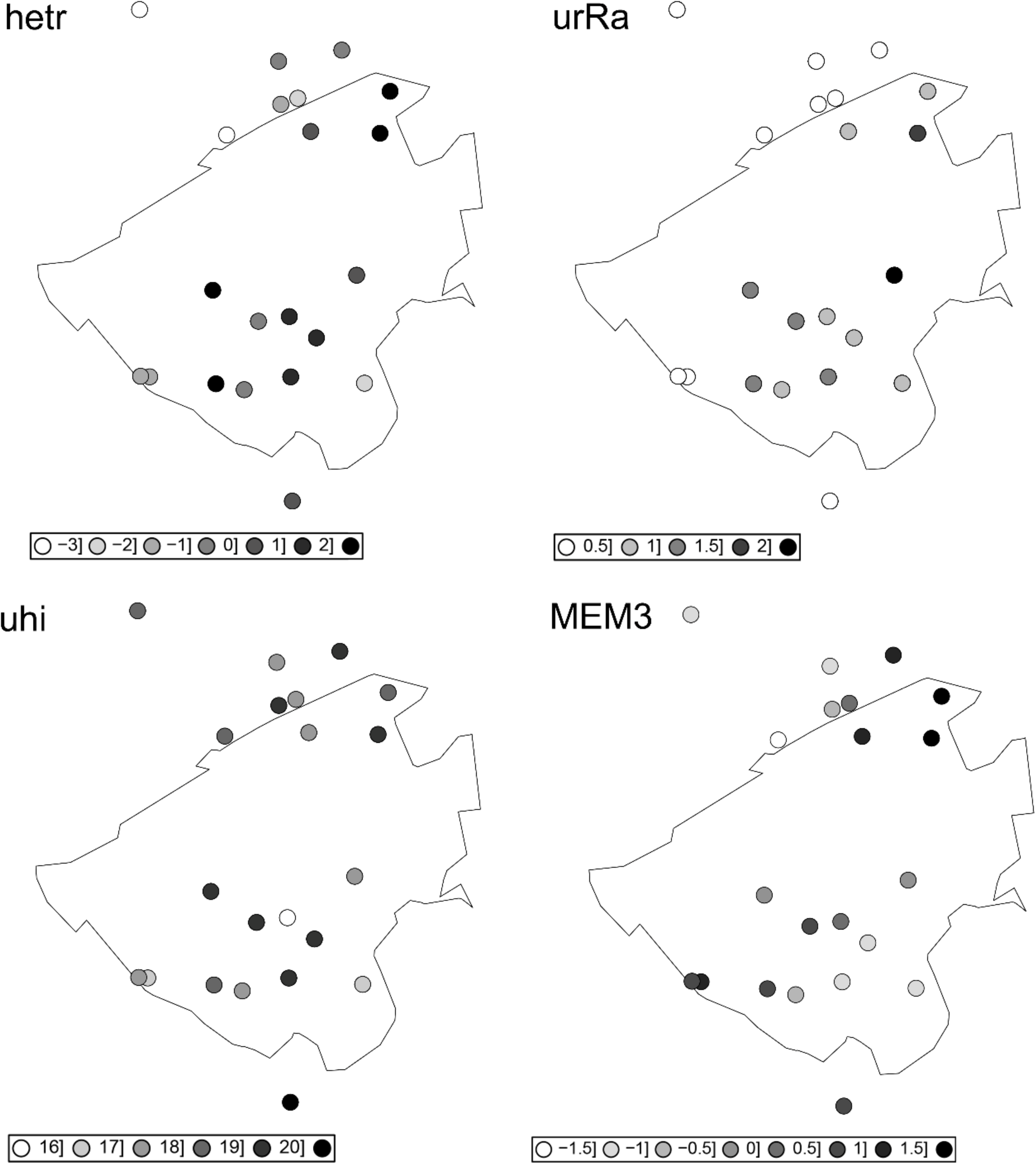
Local heterogeneity (hetr); Urbanisation rate (urRa); Urban heat island (uhi); and Moran’s Eigenvector Maps (MEM3) across urban gradient of Campo Grande. Circles represents variable values that are presented in grey gradient ranging from min to max values of each variable.

**Figure S10.**
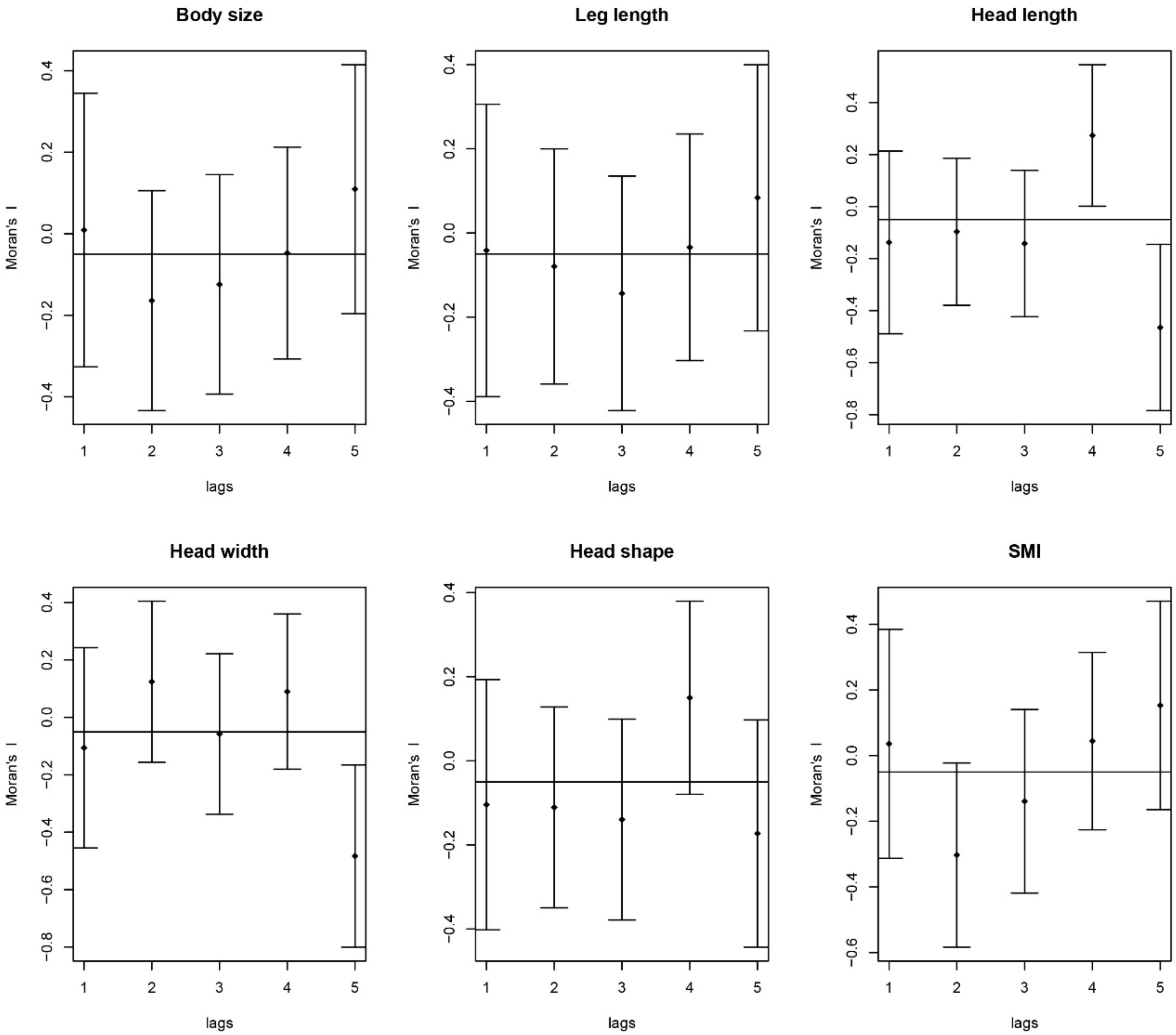
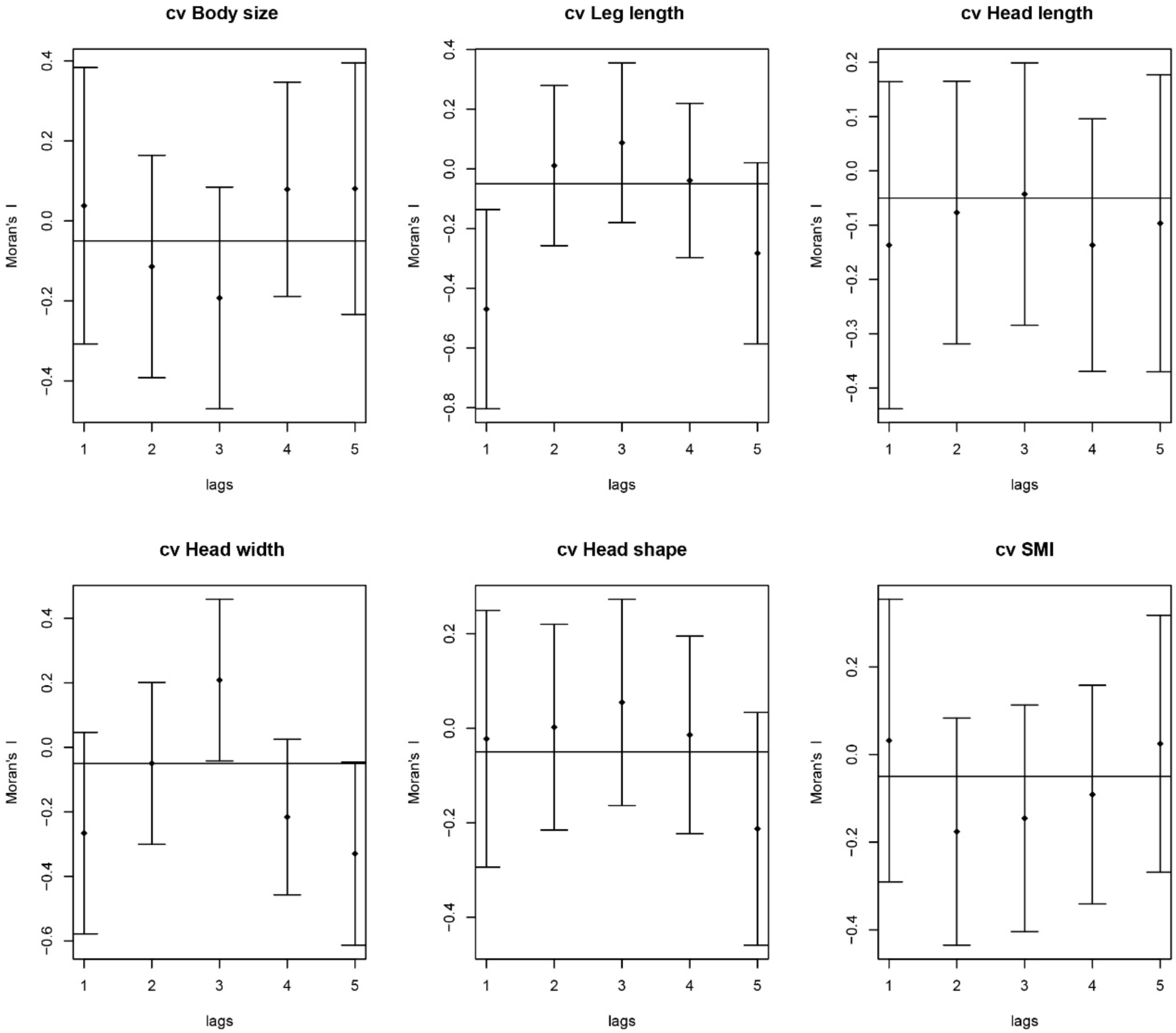
Spatial correlogram using a Moran’s *I* for the mean (this panel) and coefficient of variation (next panel) of phenotypic traits used in the Structural Equation Models. Lags are distance classes.

**Figure S11.**
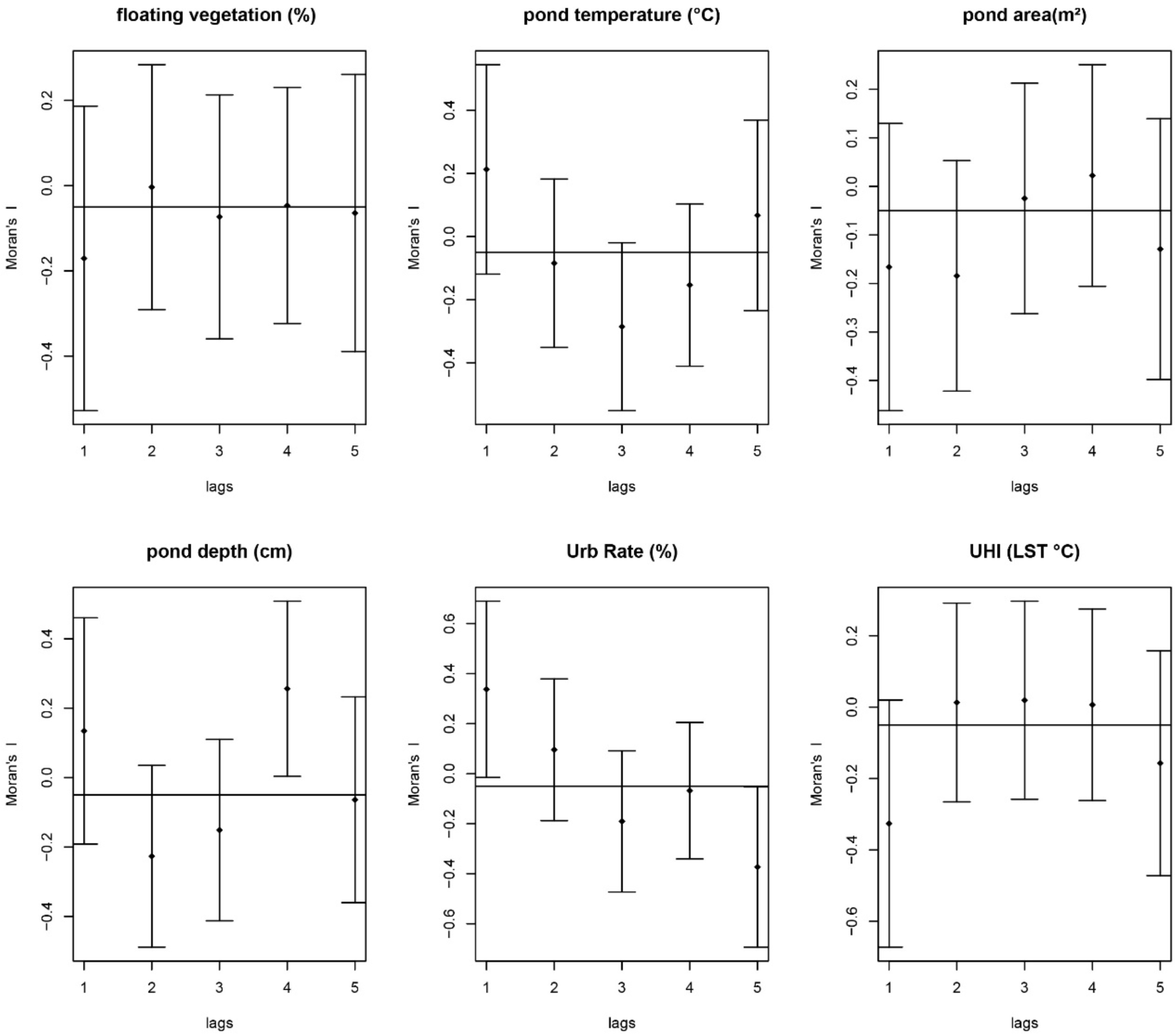
Spatial correlograms using a Moran’s *I* for local and landscape predictor variables used in the Structural Equation Models. Lags are distance classes.

**Figure S12.**
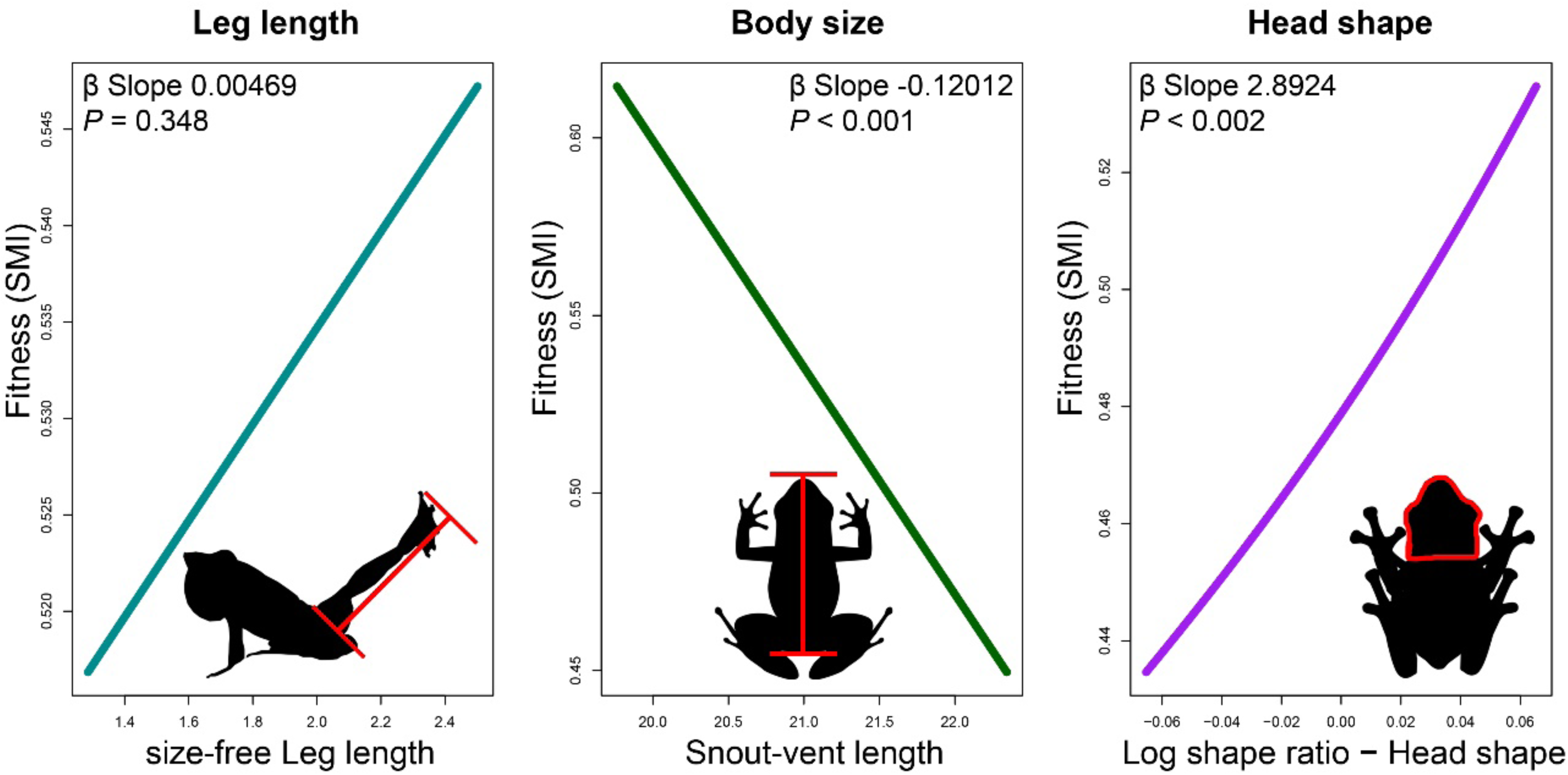
Adaptive landscape for size-free leg length (Left), body size (Centre), and head shape (log-shape ratio) (Right). Figures represent fit line of a Generalized Additive Model built using the R package gsg. The Scaled Mass Index was used as fitness proxy. Frog silhouettes are CC-BY from PhyloPic.

## Notes

https://datadryad.org/stash/dataset/doi:10.5061/dryad.3xsj3txp2

### Competing Interest Statement

The authors have declared no competing interest.

